# MDZip: Neural Compression of Molecular Dynamics Trajectories for Scalable Storage and Ensemble Reconstruction

**DOI:** 10.1101/2025.07.31.667955

**Authors:** Namindu De Silva, Alberto Perez

## Abstract

The size of molecular dynamics (MD) trajectories remains a major obstacle for data sharing, long-term storage, and ensemble analysis at scale. Existing solutions often rely on frame subsampling or reduced atom representations, which limit the utility of shared datasets. Here, we present MDZip, a neural compression framework based on convolutional autoencoders trained per system to reconstruct atomic trajectories with high geometric fidelity from compact latent representations. MDZip achieves over 95% reduction in storage size across a diverse benchmark of proteins, protein–peptide complexes, and nucleic acids. Despite operating in a physics-agnostic manner, the reconstructed trajectories accurately preserve ensemble-level features, including RMSD fluctuations, pairwise distance distributions, radius of gyration, and projections onto principal and time-lagged independent components. A residual (skip-connected) autoencoder variant consistently improves reconstruction accuracy and reduces outliers. While local structural deviations can impair energetic fidelity, short energy minimization partially recovers physically reasonable conformations. This framework enables customizable compression–accuracy trade-offs and supports a modular workflow for sharing latent representations, decoder models, and reconstruction protocols. MDZip offers a scalable solution to current storage limitations, facilitating broader dissemination of MD data without sacrificing essential dynamical information.

## 1. Introduction

Advances in computational power, longer simulation times, and growing interest in complex biomolecular systems have led to an explosion in molecular dynamics (MD) trajectory data.^1^ MD has matured into a predictive discipline, helping drive hypotheses, interpreting experiments, and determining which ligands, molecules, or protein mutants to characterize experimentally.^2–4^ Improved force fields and simulation protocols have enhanced reproducibility, and simulations are often performed in replicates to ensure robustness.

Despite these advances, MD lags behind other computational disciplines in data sharing and reusability. ^5–7^ Within academic research groups, MD data are typically generated by individual students or postdocs working on specific projects. These datasets are often stored in local directories known only to the original authors, with limited documentation or reuse by others in the lab. When researchers graduate or move on, their data can become effectively inaccessible—especially as labs run out of disk space and must prioritize newer projects. In this context, external deposition becomes the only sustainable strategy for long-term preservation and accessibility.^5,6,8^ In more recent years some academic journals and federal agencies are requesting data deposition – which is often done in free servers like zenodo that can host up to 50 gb per dataset. Since traditionally MD files take significantly larger, this often means data reduction/loss. One approach has been to store the coordinates of only some atoms (e.g., C*α* for proteins) or to downsample the frequency at which snapshots are stored. Additionally, this means storing trajectories only and rarely sharing checkpoints with coordinates and velocities that would allow for extending or reconstructing the ensemble.

One alternative to discarding data is trajectory compression. However, conventional file-level compression methods (e.g., gzip, bzip2) offer little improvement for formats already optimized for storage efficiency, such as NetCDF or DCD. A more effective strategy involves projecting trajectory data into a lower-dimensional space from which approximate reconstructions can be generated. These embeddings are highly data-efficient but inherently lossy: atomic positions in the reconstructed trajectory often deviate from their original values, and visualizations may exhibit noticeable geometric artifacts—particularly in flexible or freely rotating groups such as methyls. As a result, these approaches generally fail to preserve the resolution required for accurate local structural or energetic analysis.

For example, pcazip^9,10^ and similar algorithms apply a rotation to the coordinate system into a base of the system composed of the eigenvectors derived from the covariance matrix. These eigenvectors, in order of descending eigenvalue, capture the directions of maximum motion in the system. Thus, out of the 3*N* — 6 eigenvectors (where *N* is the number of atoms in the system), with only a few (*M << N*) of them, one can recover a large percentage of the variance of the system. Projecting the trajectory on such a set of eigenvectors will lead to a projection of each frame in the trajectory into a matrix of dimension *M* × *F*, significantly smaller than the 3*N* × *F* starting coordinates.^10^

In this work, we introduce an alternative approach using system-specific convolutional autoencoders to learn compact latent representations of MD trajectories. Our method differs from transferable, physics-based reductions in that it trains a dedicated model for each system, optimizing compression based on memorized features of the ensemble. Users can select different compression levels via the bottleneck size, balancing reconstruction fidelity and storage cost. While ensemble-level properties are accurately recovered, we observe that energy terms—being sensitive to small geometric perturbations—require a short minimization to restore physical realism. This framework enables extreme compression with minimal loss of biologically relevant structure and dynamics, offering a practical path forward for trajectory archiving, sharing, and large-scale analysis.

## 2. Methods

### 2.1. MD data

Many tools in computational structural biology and AI-based modeling have been developed and benchmarked primarily on globular proteins. To evaluate the transferability of our compression framework across a broader range of biomolecular systems, we assembled a benchmark dataset comprising seven representative targets: two proteins (insulin and the Brd3-ET domain), two protein–peptide complexes (Brd3-ET:–TP and Brd4–ET:NSD3), two 18-mer DNA duplexes with contrasting base compositions (GC-CGCGCGCGCGCGCG-GC: DNA-CG, and GC-TATATATATATATA-GC: DNA-TA), and *E. coli* Valine tRNA (Figure 1).

**Figure 1:**
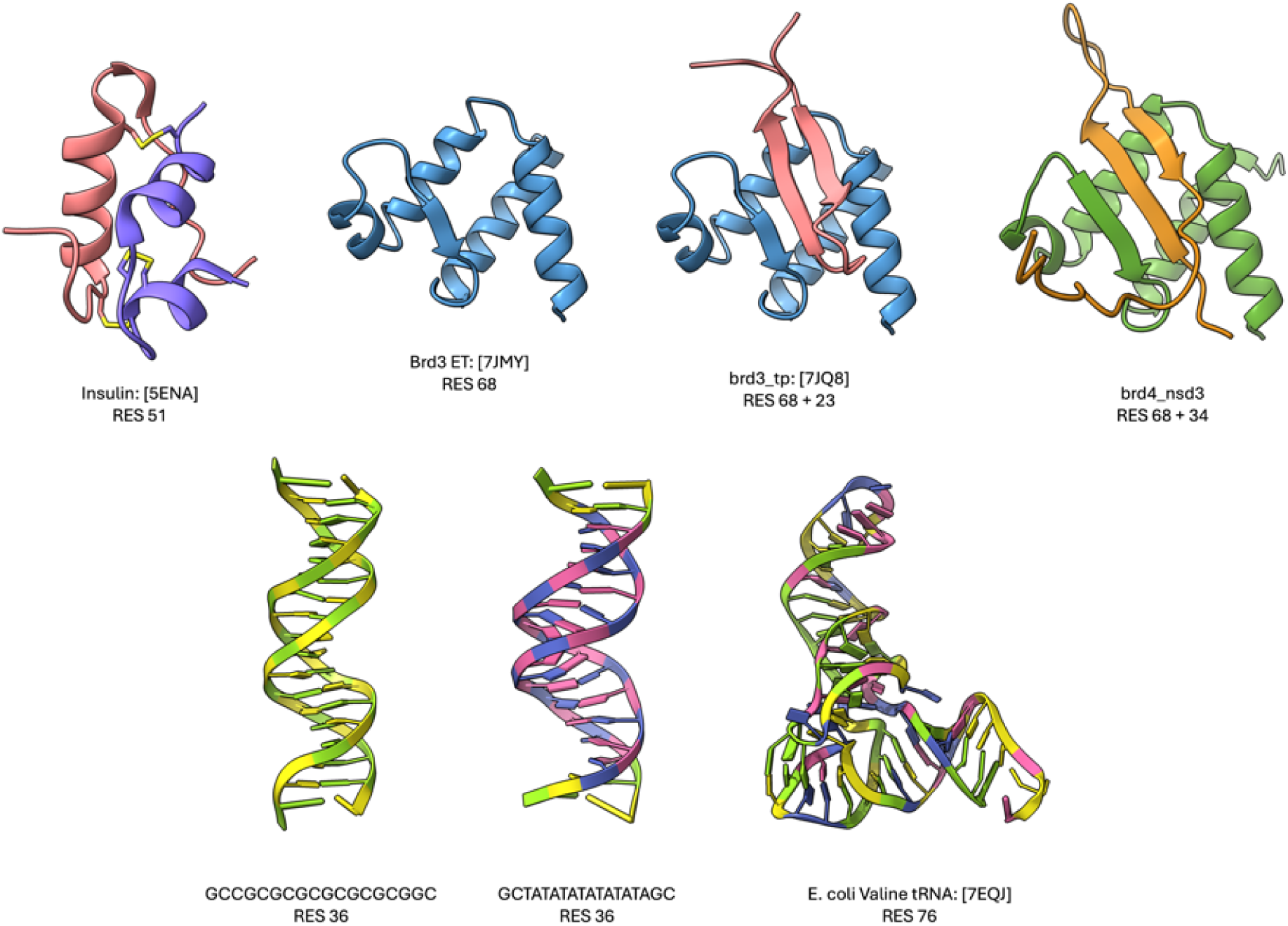
Molecular systems used to evaluate the compression algorithm. PDB IDs are provided in brackets where available, and the total number of residues is indicated as the sum across all chains in each system.

For each system we used 10^5^ frames for training, originating from either 100 ns MD saving snapshots every 1*ps* or from 1*µs* simulations retaining frames every 10*ps*. For the Brd protein systems, the first 28 N-terminal residues were removed to focus on the structured ET domain. While longer simulations are available for some of these systems, the 100 ns trajectories used here provide a diverse and standardized testbed to evaluate the robustness of our compression approach across systems with varying size, secondary structure content, and conformational flexibility.

### 2.2. Convolutional Autoencoders

We implemented two deep learning architectures to compress molecular dynamics (MD) trajectory data: a baseline convolutional autoencoder (AE) and a variant with residual skip connections (skipAE). Both models operate on individual trajectory frames, where the three-dimensional atomic coordinates are represented as an array of shape (*N*_atoms_,3). These are reshaped into a (1, *N*_atoms_,3) tensor and processed into a compact latent vector of fixed dimension *z* (see Figure 2).

**Figure 2:**
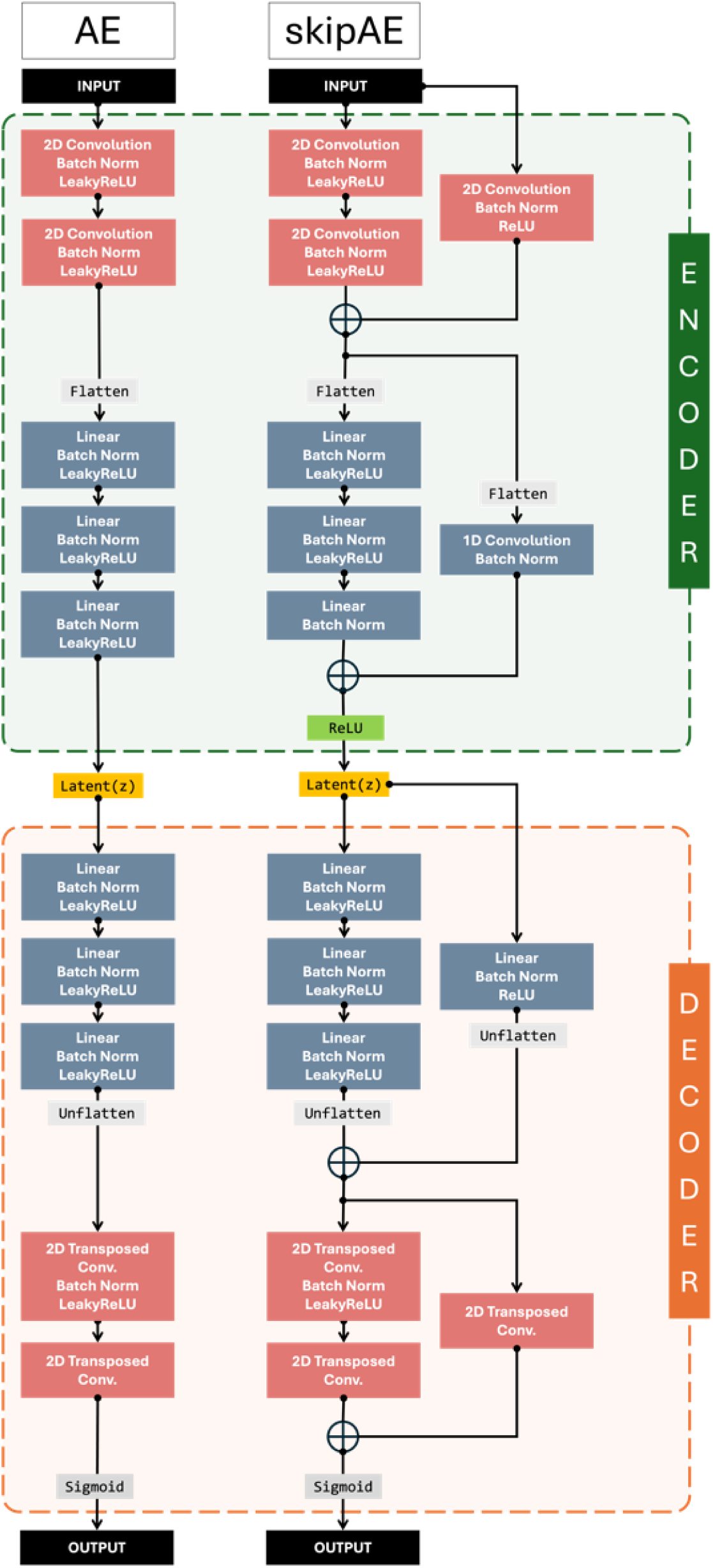
Fundamental blocks of the autoencoder architectures introduced: AE and skipAE.

The encoder starts with two 2D convolutional layers that operate over the atom dimension (as a spatial axis) and the coordinate channels (X, Y, Z). Each convolution is followed by batch normalization and LeakyReLU activation (to ensure training stability and effective gradient propagation). The output is flattened and passed through a stack of three fully connected layers to produce the latent representation (default dimension: *z* = 20). The decoder reverses this process: dense layers expand the latent space, followed by an unflattening step and two transposed convolutional layers to recover spatial structure. A final sigmoid activation maps coordinates back to the normalized [0,1] range. The network’s channel capacity scales automatically with system size, maintaining appropriate representational capacity across molecular systems of varying complexity (see SI Fig. 1).

The skipAE incorporates residual pathways to enhance information flow throughout the network architecture. While maintaining the core convolutional and fully-connected structure of the conventional model, skipAE introduces strategic bypass connections that enable direct signal propagation across multiple layers.^11^ This residual architecture addresses the vanishing gradient problem inherent in deep networks and facilitates more effective training of complex spatial relationships in molecular structures. The skip connections preserve fine-grained structural information that may otherwise be attenuated during compression, enabling the network to learn residual feature adjustments rather than complete transformations at each layer. This approach typically yields superior reconstruction fidelity and convergence properties compared to traditional autoencoders,^11–13^ proving particularly advantageous for the high-dimensional structural data characteristic of molecular trajectories.

### 2.3. Compression and Decompression

#### 2.3.1. Pre-processing MD data

Convolutional autoencoders (CAE) are particularly well-suited for molecular dynamics trajectory data due to their ability to preserve spatial relationships and learn effective feature representations through convolutional operations. ^14,15^ Since each trajectory frame contains three-dimensional atomic coordinates, MD data can be conceptualized as sequential 2D spatial data amenable to convolutional processing. Given our primary objective of trajectory compression with minimal information loss, all available positional data were utilized for training without conventional train-test partitioning, as the model must learn to reconstruct the complete dataset.

Prior to model training, several critical preprocessing steps were implemented to ensure efficient pattern recognition and learning convergence. The atomic coordinates in each trajectory frame belong to the special Euclidean group *SE*(3), where structures may undergo rotation and translation while preserving interatomic distances. To achieve *SE*(3) invariance, all trajectory frames were aligned to a reference structure (first frame) and center-of-mass coordinates were removed, ensuring that structurally similar conformations exhibit consistent positional representations.

Feature scaling represents an essential preprocessing step for ensuring numerical stability and optimal model convergence. The *SE*(3)-invariant coordinate data were normalized using min-max scaling prior to training, as described in Eq. (1). This scaling was applied independently to each spatial dimension (*X̂*, *Ŷ*, *Ẑ* coordinates) to prevent features with larger magnitudes from dominating the learning process, thereby promoting balanced representation across all spatial dimensions.

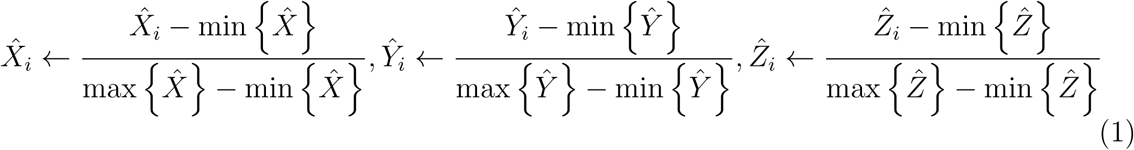

During training, trajectory frames are shuffled to prevent batch-level bias and avoid overfitting to local temporal correlations. This shuffling is applied only to the training phase of the autoencoder to ensure robust learning of the mapping between coordinates and latent representations. Importantly, compression and decompression are performed on the original time-ordered trajectory, and thus, temporal information is fully preserved in the reconstructed trajectories.

#### 2.3.2. Compression and Decompression

As outlined in the previous section, trajectory positions (*T* ϵ ℝ^*F*×*N*×3^) are preprocessed into batches of size 128 and fed to both CAE models. The models were trained with a learning rate (*α*) of 1 × 10*^—^*^4^ and weight decay (*λ*) of 0.1 across all systems using the AdamW optimizer, with all other parameters set to PyTorch default values. Unlike the standard Adam optimizer, AdamW provides the advantage of decoupling weight decay from gradient updates, which often leads to superior convergence properties.^16^ For reproducibility, the random seed was set to *42* for all compressions performed in this work.

Upon training completion, trajectory data are reloaded and preprocessed without shuffling. These data serve as inputs to the encoder of the trained CAE model with the desired latent dimensionality. The resultant latent vectors are saved as the compressed trajectory (*Z* ϵ ℝ^*F*×*l*^) alongside the corresponding trained model parameters.

During decompression, the saved latent vectors are fed to the decoder of the trained CAE model. The decoder upsamples the latent representation back to the scaled coordinate space. The same min-max linear transformer used during preprocessing is applied to map the reconstructed coordinates from the normalized space back to the original target coordinate system.

#### 2.3.3. Refining decompressed trajectories

Upon reconstruction, aberrant structures are occasionally observed due to information loss during the compression-decompression process. These small differences in pairwise RMSD between reconstructed and target structures can nonetheless lead to substantially different potential energies and structural properties. Consequently, post-reconstruction refinement procedures were implemented to address these artifacts through two complementary approaches: short energy minimization and MD steps.

Energy minimization was employed to optimize atomic positions and eliminate unfavorable contacts or overlaps. A small number of energy minimization steps were applied to each frame using the OpenMM Python module,^17^ which utilizes the L-BFGS optimizer to minimize the potential energy function. Since the primary objective was structural refinement rather than achieving global energy minima, the algorithm was implemented with early termination criteria to prevent over-optimization while ensuring the removal of geometric artifacts.

Given that most molecular dynamics simulations save snapshots at discrete intervals due to storage limitations, brief MD simulations (1*fs*) were employed as an additional refinement strategy, assuming minimal conformational changes over such short timescales. Each reconstructed snapshot was subjected to short MD simulations using appropriate AMBER force fields (ff14SB^18^ for proteins, bsc1^19^ for DNA and bsc0^20^ for the RNA system) with a Langevin integrator at 300 K temperature to allow local relaxation and improved structural consistency.

### 2.4 Analysis

#### 2.4.1 Contact maps

Contact maps provide crucial information about protein structure and folding patterns by representing pairwise Euclidean distances between atoms. These distance matrices offer insights into spatial relationships and structural contacts within protein conformations. Distance matrices were computed using the MDTraj^21^ Python package in conjunction with NumPy for numerical operations, incorporating all atoms including hydrogen atoms in the calculations. For each molecular system, we calculated mean pairwise Euclidean distances by averaging over all trajectory frames, generating representative contact maps that capture the average structural organization throughout the simulation.

#### 2.4.2 PCA and tICA

Principal Component Analysis (PCA)^22^ and Time-lagged Independent Component Analysis (TICA)^23^ are dimensionality reduction techniques essential for visualizing and characterizing the essential dynamics of molecular ensembles. Principal components were calculated using the scikit-learn^24^ Python package, with three-dimensional atomic coordinates from target trajectories, including all atoms and hydrogen atoms, serving as input for covariance matrix computation and PCA transformation. The same transformation matrix was subsequently applied to both AE and skipAE reconstructed trajectories to ensure consistent projection spaces for comparative analysis.

TICA analysis followed analogous procedures using the DeepTime^25^ Python package. A lag time of 20 ps was employed to capture slow conformational transitions characteristic of biomolecular dynamics. The TICA transformation was fitted to target trajectory coordinates and then applied to predicted trajectories from both autoencoder models, enabling direct comparison of dynamical projections and assessment of essential motion preservation during compression and reconstruction.

#### 2.4.3 Energy analysis

Energy calculations were performed using cpptraj,^26^ the integrated trajectory analysis program within the AMBER^27^ molecular dynamics suite. All trajectory frames, including hydrogen atoms, were analyzed to compute six distinct energy components: bond, angle, dihedral, electrostatic, van der Waals, and total energies. Electrostatic interactions were calculated using the Particle Mesh Ewald (PME)^28^ methodology with a direct space cutoff of 8 Å, representing the standard default parameter for accurate long-range electrostatic treatment in biomolecular simulations. All force field parameters required for energy calculations were extracted from system-specific topology files, with identical topology files used consistently across target, AE, and skipAE trajectories to ensure comparable energy evaluations. This approach enabled direct assessment of energetic fidelity between original and reconstructed molecular conformations.

#### 2.4.4 DNA helical parameters

Nucleic acid helical parameters were calculated using cpptraj following the Tsukuba convention as implemented in 3DNA.^29,30^ Base-pair axes were defined with a distance cutoff of 2.5 Å between base-pair axis origins to determine eligible base-pairing interactions. To ensure consistent base-pair and base-pair step definitions across target, AE, and skipAE trajectories, a reference structure was selected from the target trajectory exhibiting well-preserved base-pairs with minimal structural distortion.

Base-pair parameters (shear, stretch, stagger, buckle, propeller, opening) and base-pair step parameters (shift, slide, rise, tilt, roll, twist) were computed for all trajectory frames. During comparative analysis between target and predicted structures, frames containing null values arising from base-pair disruption beyond the distance cutoff were excluded from statistical calculations to maintain data integrity and ensure meaningful structural comparisons.

Force constant analysis^31^ was performed using the calculated base-pair and base-pair step parameters to assess the flexibility and constraints of nucleic acid structural features. Force matrices were derived by inverting the covariance matrices calculated separately from base-pair parameter data and base-pair step parameter data. This inversion procedure yields force constant matrices where higher values indicate lower structural variance, analogous to stronger harmonic restraints in the respective degrees of freedom. All matrix calculations and inversions were performed using the NumPy Python library, providing quantitative measures of structural rigidity and flexibility for comparative analysis between target and reconstructed nucleic acid conformations.

## 3. Results and discussion

### MDZip achieves vast compression with accurate ensemble reconstruction

Training convergence was achieved within 1000 epochs for all molecular systems, with loss curves exhibiting stable plateau behavior (SI Figure 2). The absence of oscillatory or erratic fluctuations indicates that the optimization process successfully reached adequate performance levels across the entire benchmark dataset. Importantly, this consistent behavior was observed across both protein and nucleic acid systems, suggesting that the performance of the algorithm is not strongly dependent on the type of biomolecule being studied. This highlights the robustness of the training protocol and the transferability of the chosen hyperparameters across a broad range of structural complexities.

The compression ratio is dictated by the user-defined bottleneck dimensionality, with each molecular dynamics snapshot compressed into the corresponding latent vector representation. Using a default bottleneck dimension of 20, we achieved over 95% compression across all molecular systems (Figure 3A). Since both AE and skipAE models employ identical bottleneck dimensions, compression ratios remain consistent between architectures. As all the systems contain the same number of frames and use the same bottleneck size, the compressed trajectories are uniform (37.3 MB in this set). Notably, trajectories with higher atom counts exhibit proportionally greater compression efficiency. It is also possible to specify a compression value, which will automatically calculate the size of the latent space (*z*).

**Figure 3:**
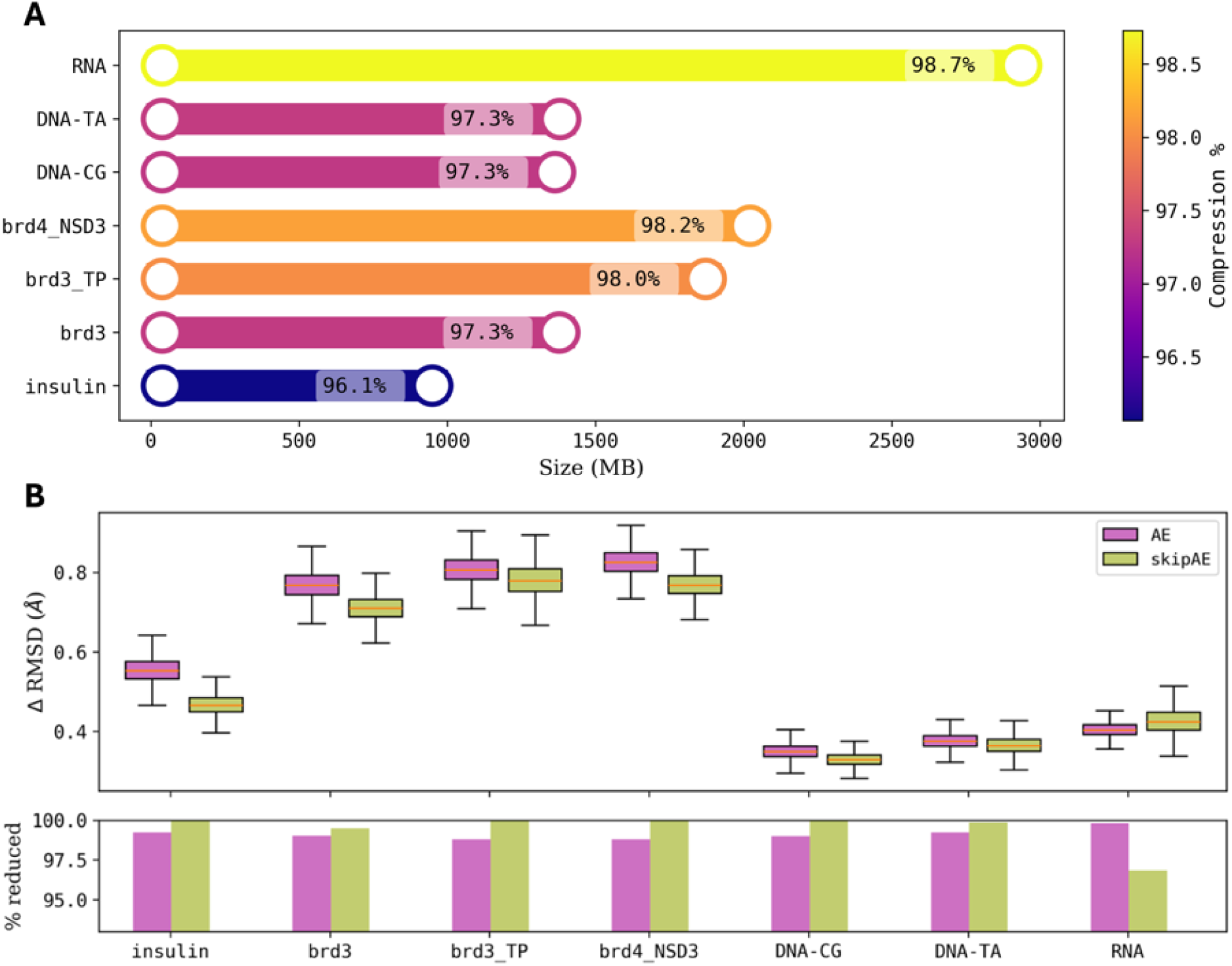
Compression performance and reconstruction fidelity across molecular systems. A A. Barbell plots show original (right) and compressed (left) trajectory sizes, with compression percentages labeled. B Reconstruction accuracy assessment: (top) pairwise RMSD (Δ*RMSD*) between original and reconstructed structures shown as box plots comparing conventional autoencoder (AE, magenta) and skip-connection enhanced autoencoder (skipAE, green), with median values indicated by orange lines (Lower RMSD values indicate better reconstruction fidelity.); (bottom) percentage of structures reconstructed with Δ*RMSD* below 1 *Å* for protein systems and below 0.5 *Å* for nucleic acid systems.

Current open-access repositories such as Zenodo^32^ impose storage limitations of 50 GB per record with a maximum of 100 files. However, contemporary molecular simulation studies increasingly generate large-scale trajectories extending to mesoscopic scales, enabled by advancing computational capabilities and methodological development. ^33–35^ Moreover, individual studies often produce multiple such trajectories, exacerbating storage constraints. Conventional approaches to trajectory size reduction include solvent stripping or frame subsampling with temporal strides. However, achieving compression levels comparable to our method would require the removal of approximately 95% of trajectory frames, resulting in substantial data loss and rendering trajectories scientifically inadequate. As an example, A typical MD project comprising three protein variants, each simulated in triplicate for 10*µs* and saved every 1 ps, produces ∼ 10 million frames per trajectory—an uncompressed total of ∼ 671 GB (dry trajectories) across the nine trajectories. Our AI-based compressor (latent space z=20) shrinks this dataset by 95 %, delivering the full, all-atom, every-frame ensemble in just 33.6 GB, well below Zenodo’s 50 GB record limit. Conventional strategies might use other trade-offs: keeping all atoms but subsampling 1 frame in 100 lowers the dataset to 6.7 GB yet discards 99 % of temporal information; stripping to *C*_α_ atoms retains time resolution but still weighs in at 34.9 GB and loses side-chain and solvent detail; even the extreme combination of *C*_α_-only plus 1 : 100 subsampling reaches a tiny 0.35 GB, but at the cost of both temporal and structural fidelity.

Remarkably, these AI compression levels are achieved while maintaining high structural fidelity, with reconstructed conformations closely resembling target structures and exhibiting mean pairwise root mean squared deviations (Δ*RMSD*) below 1*Å* (Figure 3B). The skip-connection enhanced autoencoder demonstrates superior performance compared to the conventional autoencoder, producing structures with lower median Δ*RMSD* across all systems except RNA, highlighting the model’s enhanced reconstruction capabilities. We noticed that, 1000 epochs with more learnable parameters induce overfitting, and that for simpler dynamics (lower motion) using simpler models is better (DNA and RNA).

Both architectures occasionally generate outlier predictions with Δ*RMSD* exceeding 1*Å*, though skipAE consistently produces fewer such instances (Figure 3B(bottom)). Specifically, skipAE achieves exceptional performance with zero outliers above 1*Å* for four systems (insulin, DNA-CG, DNA-TA, and RNA) and minimal outlier rates for the remaining systems (0.51% for Brd3). In comparison, the conventional AE exhibits higher but still remarkably low outlier percentages above 1*Å*: 0.775% (insulin), 0.985% (Brd3), 1.212% (Brd3 TP),1.194% (Brd4 NSD3), 0% (DNA-CG), 0.023% (DNA-TA), and 0% (RNA) (notice that outlier limits in Figure 3B use the more stringent 0.5*Å* threshold for nucleic acids). Collectively, outlier frames constitute less than 2% of the total dataset across all systems, representing a better alternative to describing the ensemble than conventional stride-based trajectory reduction approaches that would require removal of ∼ 95% of frames to achieve comparable compression ratios.

### MDZip preserves global properties of the ensemble

To assess the quality of reconstructed trajectories, we evaluated several ensemble properties that characterize global structural behavior. RMSD represents the most fundamental metric for quantifying ensemble structural variations. We analyzed RMSD fluctuations using the closest frame to the average geometric positions of the atoms (average structure) of each trajectory as the reference structure. As shown in Figure 4A, the RMSD of reconstructed trajectories maps on top of the original RMSD across all systems.

**Figure 4:**
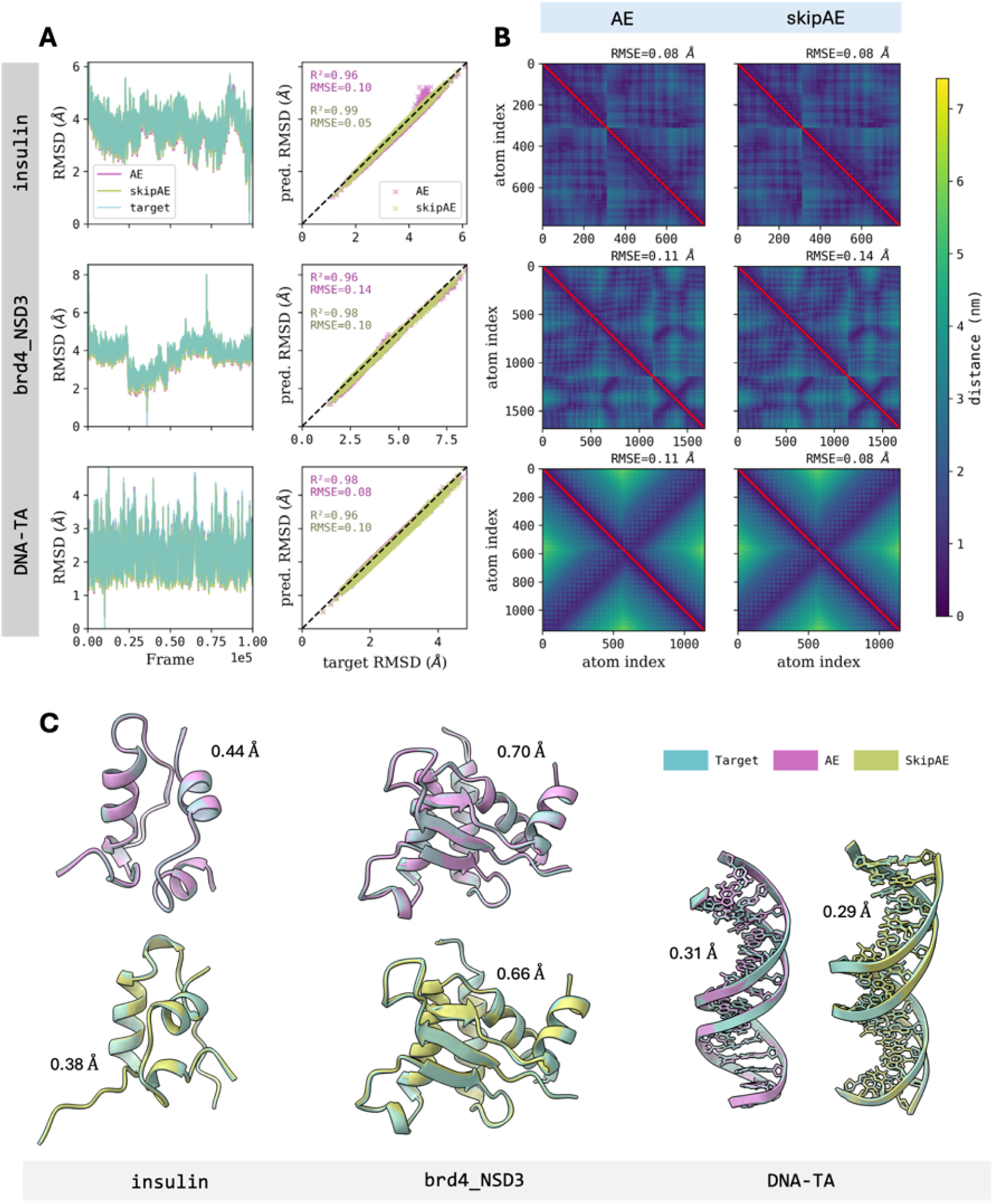
Reconstruction performance analysis across representative molecular systems. (A) RMSD trajectory fluctuations and correlation plots for target (cyan), AE (magenta), and skipAE (green) with *R*^2^ and RMSE values. (B) Distance matrices showing target (upper triangle) and predicted (lower triangle) distances with RMSE values. (C) Best reconstructed structures with Δ *RMSD* values displayed. Complete analysis for all systems in SI Fig. 3.

Both autoencoder models demonstrate strong correlation with target trajectories, achieving high *R*^2^ values for most systems when evaluated against the identity line (y=x). Notably, the protein-ligand complexes (Brd3 TP and Brd4 NSD3) show slightly reduced correlation coefficients, likely attributable to the additional conformational complexity introduced by ligand binding dynamics. Root mean square error (RMSE) values follow similar trends, with skipAE consistently outperforming the conventional model across all systems except RNA, maintaining reconstruction errors below 0.2 *Å*. The DNA-TA system exhibits the lowest RMSE of 0.07 *Å*, demonstrating exceptional reconstruction fidelity using skipAE. Interestingly, for larger RMSD values, AE is slightly misbehaved for some systems, whereas the skipAE corrects for these issues.

To further assess structural preservation, we analyzed pairwise distance matrices (contact maps), which provide critical insights into contact preservation—particularly important for protein folding and stability. Distance matrix analysis reveals excellent reconstruction accuracy with RMSE values ranging from 0.07 — 0.16 *Å* for all the molecular systems. The mean distance matrices (Figure 4B) appear as nearly perfect mirror images along the diagonal, indicating preservation of intramolecular contacts. These comprehensive analyses demonstrate that both architectures successfully capture global structural dynamics and local contact patterns.

Principal Component Analysis (PCA) and Time-lagged Independent Component Analysis (tICA) provide complementary approaches for capturing global ensemble dynamics by projecting high-dimensional coordinate data into interpretable lower-dimensional representations. To evaluate whether our autoencoder models preserve essential dynamical features, we analyzed the first two principal components (PC1 vs PC2) and the first two time-lagged independent components (TIC1 vs TIC2) for each molecular system (Figure 5).

**Figure 5:**
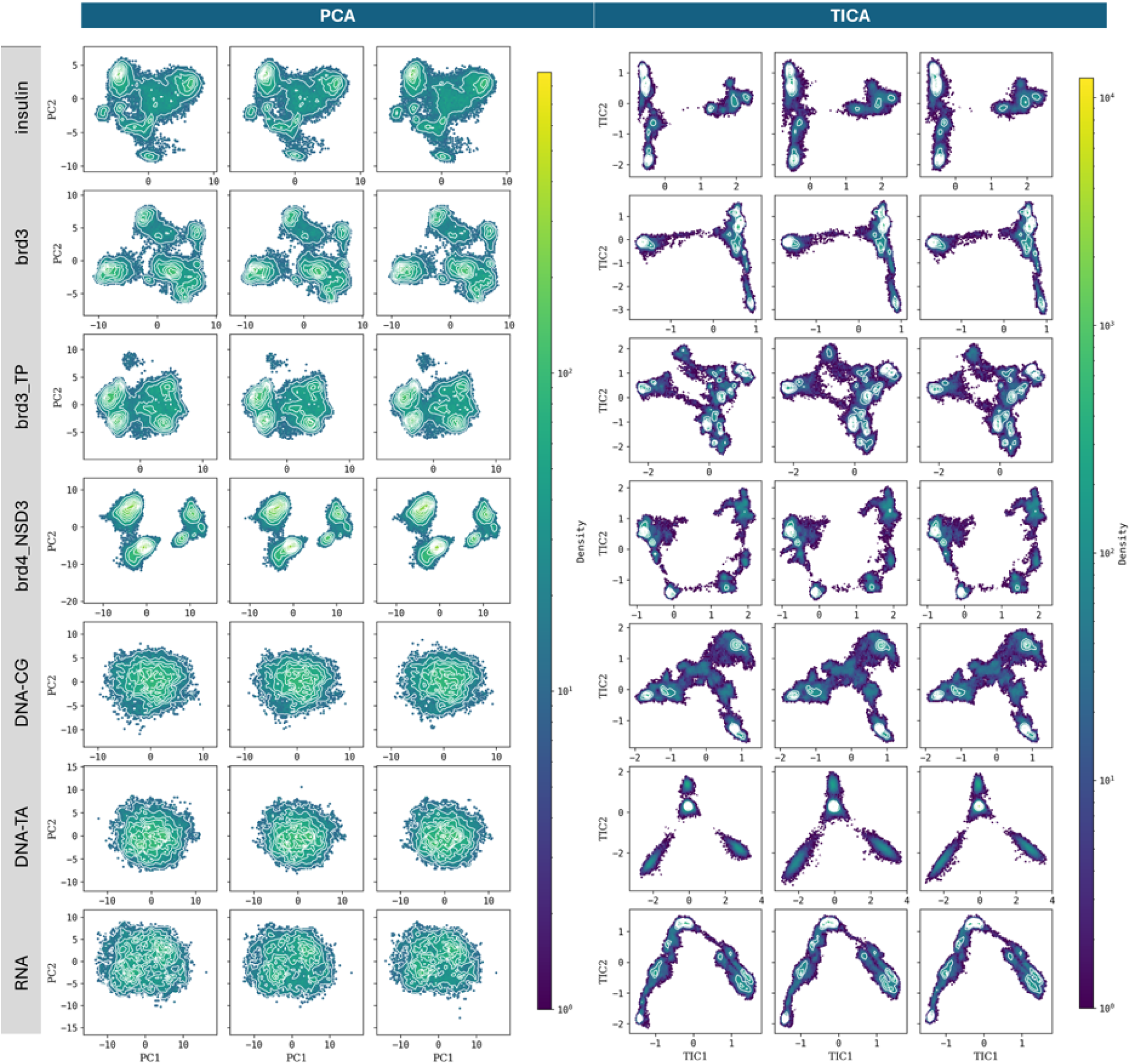
Principal Component Analysis (PCA) and Time-lagged Independent Component Analysis (tICA) projections for target and reconstructed trajectories across molecular systems. Each row represents a distinct molecular system. (Columns 1-3) Projection on two first PCA eigenvectors (PC2 vs PC1) of original, AE, and skipAE ensembles (columns 1,2, and 3 respectively). (Columns 4-6) tICA projections (TIC2 vs TIC1) with 20 ps lag time for target trajectories (column 4), AE predictions (column 5), and skipAE predictions (column 6). Both models successfully preserve the essential dynamical features and conformational landscapes of the original trajectories.

tICA analysis was performed using a lag time of 20 *ps* to capture slow conformational transitions characteristic of biomolecular dynamics. Both autoencoder architectures successfully reproduced the target trajectory projections in both PCA and tICA space, demonstrating their ability to preserve essential dynamical information during compression and reconstruction. Notably, the similarity between target and reconstructed tICA projections confirms that temporal correlations are maintained. This is consistent with the fact that only the training phase involves shuffled frames, while compression and reconstruction are performed on the original time series, preserving the trajectory’s dynamical structure.

Quantitative assessment reveals near-perfect correlation between target and reconstructed projections, with coefficient of determination values approaching unity (*R*^2^ ∼ 1) for all principal and independent components (SI Figure 4). The *R*^2^ values were calculated using identity line fitting (*y* = *x*), representing the proportion of variance in reconstructed projections explained by the target trajectory dynamics. These results confirm that both models effectively maintain the global conformational landscape and dynamical behavior of the original trajectories, validating their utility for preserving biologically relevant structural information despite extreme compression ratios.

### Limitations and opportunities in local reconstruction

Lossy compression inherently involves trade-offs between preserving global ensemble behavior and retaining local structural details such as bond lengths, dihedrals, and atomic contacts. To assess local fidelity, we analyzed bond distance preservation across all molecular systems. Protein systems showed mean bond deviations of ∼0.2 Å, while nucleic acid systems exhibited tighter preservation (0.05 Å), particularly in base-paired regions (Figure 6A). The skip-connection enhanced autoencoder consistently outperformed the baseline AE across all systems.

**Figure 6:**
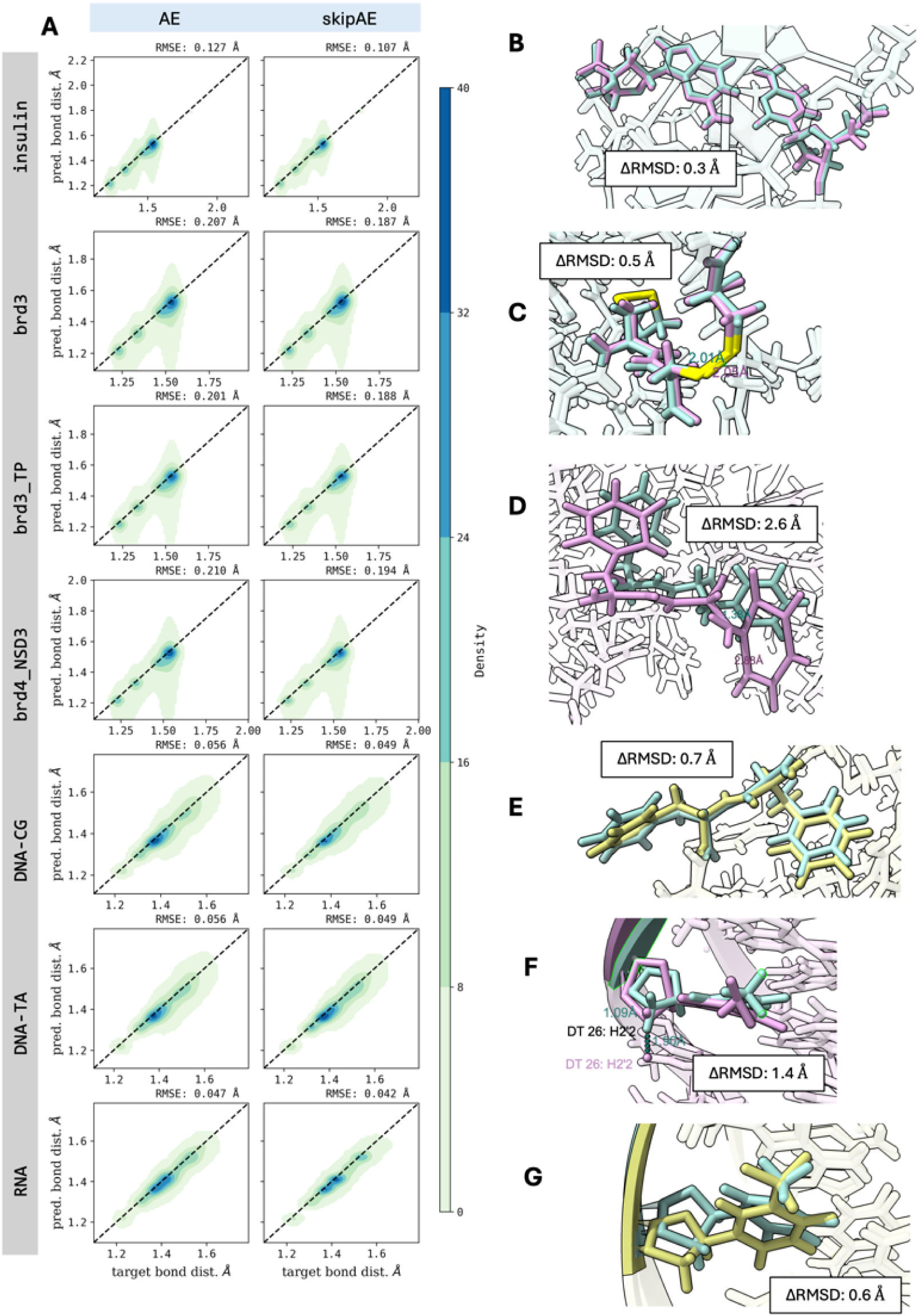
Bond distance preservation and structural feature reconstruction quality. (A) Contour plots of predicted versus target bond distances for AE (column 1) and skipAE (column 2) with RMSE values displayed. (B-G) Representative structural features showing nucleotide base rings, disulfide bonds, and examples from high-error frames for phenylalanine rings and thymine bases. Color coding: target (blue), AE (magenta), skipAE (green). ΔRMSD values shown in boxes for each structure.

While average bond deviations remained low, occasional outliers were observed, with deviations up to ∼1.5 Å for protein systems and ∼1 Å for nucleic acid systems. Analysis reveals that model performance correlates strongly with overall reconstruction quality: structures with low Δ*RMSD* values exhibit excellent preservation of critical structural features, including carbon ring geometries and disulfide bond distances (Figure 6B,C). Conversely, structures with high ΔRMSD values show proportionally degraded local structural quality (Figure 6D-G). A common issue involves hydrogen-involving bonds, which in MD are constrained using SHAKE or similar algorithms.^36,37^ These produce discrete distance distributions that are not explicitly learned in our geometry-based models, contributing to larger reconstruction errors in constrained degrees of freedom.

Further analyses are grouped by biomolecule type to accommodate structural metrics unique to each class.

### Protein and protein-peptide

For protein systems, we evaluated backbone dihedral angles (*ϕ*, *ψ*), radius of gyration (Rg), and secondary structure using the Dictionary of Secondary Structure of Proteins (DSSP).^38^ For protein–peptide complexes, we also analyzed the distance between the centers of mass (COM) of the protein and peptide components to assess binding geometry preservation.

Ramachandran plots provide a sensitive structural fingerprint of protein conformations, capturing backbone dihedral angle distributions and secondary structure preferences. Both autoencoder models closely reproduced the two-dimensional distributions of Ф and Ψ angles observed in the target structures (Figure 7A,B and SI Fig 5). This qualitative agreement is supported quantitatively by dihedral angle reconstruction errors in the 10–15° range, indicating robust preservation of local backbone geometry.

**Figure 7:**
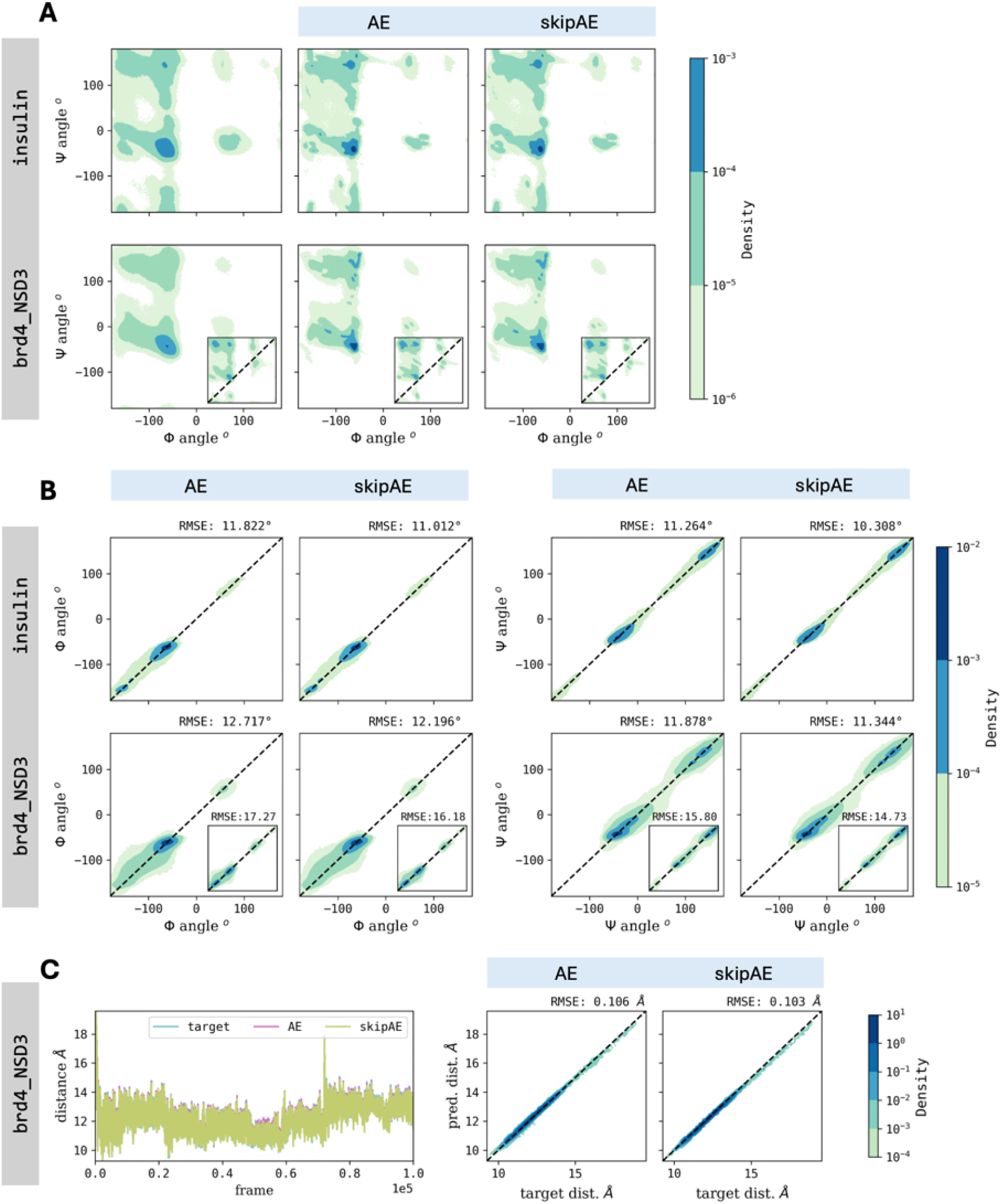
Local analysis for protein systems (A) Ψ versus Ф angle distributions for original, AE, and skipAE. (B) Ф and Ψ correlation analysis (original vs model) (D) Center-of-mass distance analysis for protein-peptide complexes showing trajectory fluctuations and correlations. Peptide data shown in the inset plots where applicable. RMSE values are displayed at the top right. Complete analysis in SI Fig. 5.

Analysis of the radius of gyration further confirmed the models’ ability to preserve global compactness, with reconstruction errors of approximately 0.1*Å* across all protein systems (see SI Figure 6). The conventional AE model produced more outlier predictions than the skipAE, as evidenced by consistently higher RMSE values, highlighting the enhanced accuracy of the skip-connection architecture.

Secondary structure was evaluated using DSSP, which assigns elements based on hydrogen bonding patterns. Both models successfully recovered the secondary structure profiles of the target trajectories, preserving key helical and *β*-sheet regions across all protein systems (see SI Figure 7). Peptides in the protein–peptide complexes also exhibited high-fidelity reconstruction, with *β*-structures retained throughout. Together, these results demonstrate that both autoencoder models preserve critical features from local backbone geometry to higher-order structural organization, with skipAE offering improved robustness.

For protein-peptide complexes, we also assessed intermolecular binding geometry preservation by measuring center-of-mass (COM) distances between protein and peptide components. Both autoencoder models achieve excellent performance with RMSE values of 0.1 — 0.2*Å*, indicating faithful preservation of binding interface integrity and overall complex architecture (Figure 7C, SI Fig 5C). This performance confirms that MDZip retains biologically meaningful binding conformations essential for protein–peptide interaction studies.

### Nucleic Acids

For double-stranded nucleic acid systems, we evaluated structural reconstruction using the standard helical parameterization of base pairs (BP) and base-pair steps (BPS), which includes three translational and three rotational degrees of freedom for each level. These descriptors rely on defining a local reference frame for each base or base pair using an origin and three orthogonal axes. BP parameters are derived from the frames of paired bases, while BPS parameters are computed from adjacent base-pair frames. Because these definitions depend on the precise positioning and orientation of multiple atoms, small deviations in atomic coordinates can propagate into larger errors in helical parameter values. To ensure consistent base-pair identification across frames and avoid misalignment artifacts, we used the first frame of each trajectory as the structural reference for all subsequent analyses.

Reconstruction of base-pair parameters showed excellent agreement for translational components. Shear exhibited RMSE values of 0.2 — 0.5*Å*, stretch showed 0.1 — 0.2*Å* error, and stagger had slightly higher deviations (0.3 — 0.5*Å*). Angular parameters were reconstructed with RMSE values between 3 — 7 for DNA systems, with opening being the most accurate (3), followed by buckle and propeller. RNA posed greater challenges, particularly for buckle parameters with RMSE values reaching ∼ 16. Propeller reconstruction in RNA achieved similar accuracy to DNA (6), while opening parameters showed approximately double the error of DNA systems (∼ 6 vs ∼ 3 in DNA) (SI Fig 8).

**Figure 8:**
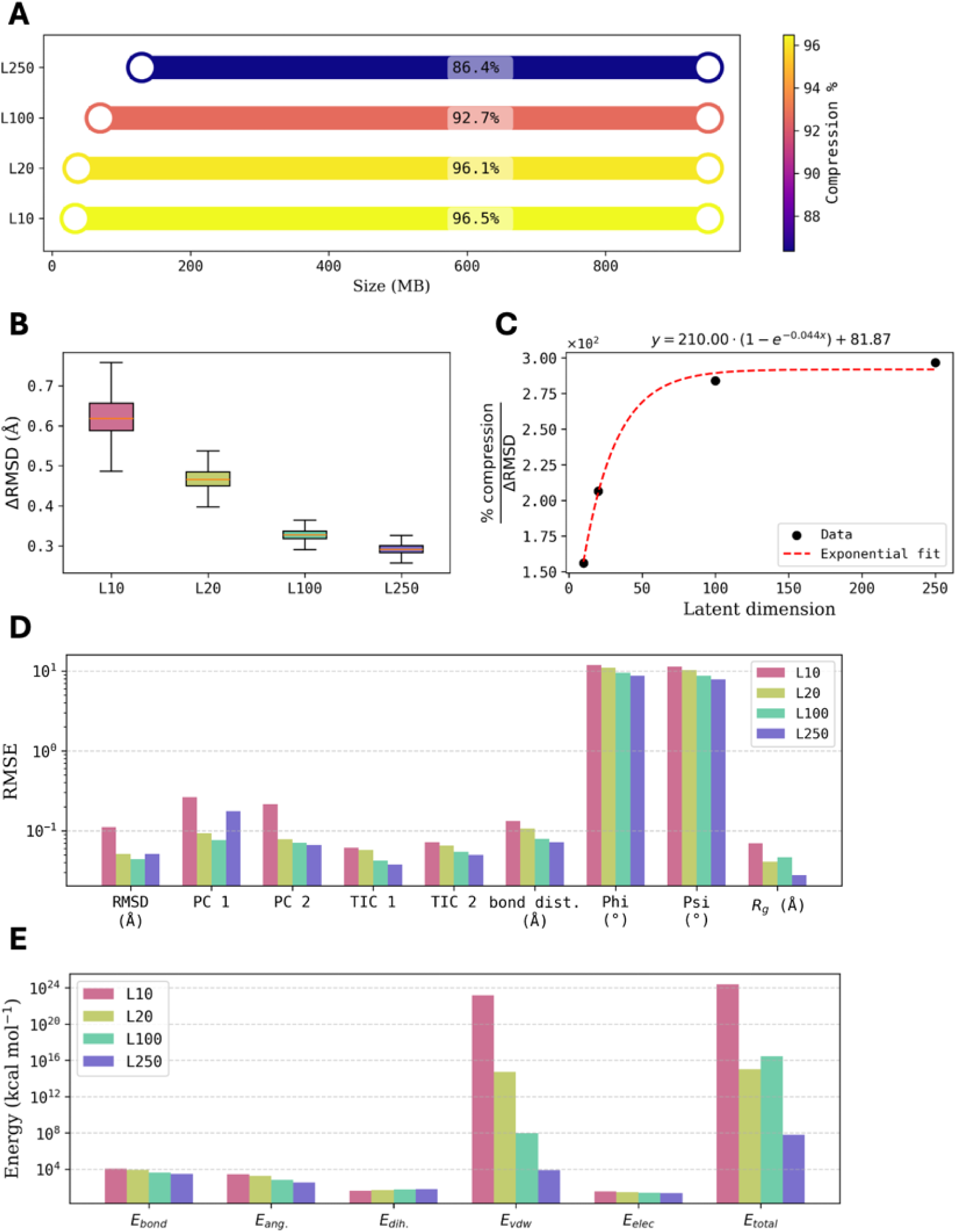
Compression model performance across bottleneck dimensions.(A) Compression ratios shown as barbell plots with initial (right) and compressed (left) file sizes. (B) Reconstruction quality (Δ*RMSD*) distributions by bottleneck size. (C) Performance optimization: compression percentage to Δ*RMSD* ratio vs. latent dimension. (D) RMSE for structural properties across bottleneck sizes. (E) Energy reconstruction accuracy by dimension. Colors indicate latent dimensions: pink (L10), green (L20), mint-green (L100), purple (L250), illustrating the compression-fidelity trade-off. (see SI Fig. 19-21 for detailed correlation graphs

Base-pair step parameters demonstrated higher fidelity compared to base-pair parameters. Translational parameters (shift, slide, rise) were reproduced with RMSE values of 0.2 — 0.3*Å*, while angular parameters (tilt, roll, twist) achieved errors between 2 — 5.5 (SI Fig. 9), highlighting the model’s ability to retain global helical features such as base stacking and twist.

To further probe local dynamical accuracy, we evaluated effective force constants by inverting covariance matrices derived from BP and BPS fluctuations (see Methods). This analysis revealed systematic biases in both models (SI Figure 10). The conventional AE model consistently overpredicted force constants, while skipAE underpredicted some translational force constants while overestimating other base-pair step force constants. Distribution analysis indicates that while mean structural properties are well preserved, variance patterns differ from target trajectories, as seen from the tails of distributions for these parameters in SI Figure 11. This suggests that the models primarily learn average structural features while constraining conformational flexibility, likely due to insufficient sampling of rare conformational states in the training data. Base-pair step parameters showed improved force constant reconstruction compared to base-pair parameters, though systematic deviations persist, highlighting the importance of comprehensive conformational sampling for optimal reconstruction quality.

### Energies

Our autoencoder framework is purely geometry-based, it does not incorporate physics-based constraints or force field information. Both models are trained to minimize coordinate deviations between predicted and target atomic positions. However, molecular energies are non-linear functions of atomic coordinates involving higher-order terms (e.g., powers, exponentials, and inverse distances), which makes them highly sensitive to even small positional errors. Consequently, geometric accuracy does not always translate into energy accuracy.

As anticipated, both models struggle to reproduce accurate energetic profiles across molecular systems (SI Figure 12-14). Energy reconstruction errors are generally larger for protein systems than for nucleic acids, except in dihedral energy terms, where protein performance improves (see SI Fig. 12-14). When only the the structures with lower reconstruction error is selected (Δ*RMSD <* 1*Å* for proteins and Δ*RMSD <* 0.5*Å* for DNA*/*RNA) we see a very significant change in dihedral energies for proteins and a visible change in total energy for DNA, mainly due to improved van der Waals (vdW) interactions. These thresholds are adjustable and depend on the user’s tolerance for outlier exclusion. In our benchmarks, applying these cutoffs retained more than 96% of the data (Figure 3B (bottom).

The systematic energy reconstruction limitations reflect the fundamental challenge of preserving non-linear molecular properties through purely geometric learning approaches. However, the skip-connection enhanced architecture demonstrates superior performance to the conventional model across all energy metrics, suggesting potential for improvement through architectural advances. These results highlight both the current limitations and future opportunities for incorporating physics-aware constraints into trajectory compression frameworks. We tested physics-based post-processing on reconstructed structures to explore mitigation strategies using energy minimization. We selected insulin as a test case due to its moderate size and limited computational demands. Minimizations were performed in OpenMM using the Amber ff14SB force field and the L-BFGS optimizer, applying harmonic restraints to preserve structural integrity. Since energy is highly sensitive to small geometry changes, we reasoned that short minimizations that remove large vdW penalties while retaining the reconstructed geometry would be best. We systematically tested protocols with 10, 50, 100, and 500 minimization steps. We also tested adding a few MD steps.

As anticipated, increasing minimization steps progressively increased structural drift, leading to higher Δ*RMSD* relative to target conformations (SI Fig. 15-17). This also worsened global ensemble properties like radius of gyration and backbone angles, particularly beyond 10 steps. Notably, MD simulations following minimization occasionally produced outlier structures, particularly when initial conformations contained poor contacts, resulting in system instability and elevated TIC2 RMSE values reaching 1.8*Å*.

Nonetheless, minimal minimization (≤ 10 steps) yielded meaningful improvements. Bond length RMSE by improved by 0.03 — 0.04*Å* from the initial 0.1*Å* baseline. Dihedral angle RMSE values of ∼ 10.9 and ∼ 10.2 for Ф and Ψ angles, respectively and low radius of gyration error (0.036*Å*) were best at 10 steps. MD simulations consistently degraded all structural metrics, highlighting the destabilizing effects of excessive optimization. Energy reconstruction improved most significantly for van der Waals terms, contributing to reduced total energy error. Electrostatic energies, however, showed minimal improvement or even degradation.

Together, these findings suggest that brief energy minimization can enhance energetic consistency without compromising structural fidelity—provided the protocol remains conservative. In contrast, extensive minimization or follow-up MD can distort ensemble properties, reducing the scientific value of compressed trajectories. As such, minimal post-processing offers a practical compromise for applications where energy accuracy is critical.

### Compression-performance trade-offs and implementation guidance

Our compression framework is designed to be universally applicable across molecular systems – and specifically trained for each ensemble. Users can train custom models on their own trajectories and adjust the level of compression by selecting an appropriate bottleneck dimension. Larger latent spaces retain more atomic detail but reduce compression efficiency, while smaller latent dimensions maximize compression at the cost of reconstruction fidelity (Figure 8). This design allows users to balance compression and accuracy based on the specific requirements of their application.

To systematically evaluate this trade-off, we tested bottleneck dimensions of 10, 20 (default), 100, and 250 using the skipAE model. As expected, reconstruction quality improved with larger latent dimensions, with diminishing returns beyond 100 dimensions. We introduced a practical performance metric—compression percentage divided by ΔRMSD—to help quantify this trade-off. This ratio increased steeply at first and then plateaued (Figure 8C), indicating that moderate bottlenecks often offer the best efficiency-to-accuracy balance.

Structural properties showed modest improvements with increasing latent size. For example, bond distance RMSE improved from 0.133 Å to 0.072 Å, dihedral angle errors improved by ∼3°, and radius of gyration RMSE decreased from 0.070 Å to 0.028 Å (Figure 8D). Energy reconstruction followed a similar trend, with the most pronounced gains in van der Waals terms, leading to lower total energy errors (Figure 8E). These results define the upper bound of what is achievable through geometry-based models. However, these modest improvements in local and global structural properties may not justify the substantial reduction in compression efficiency for many applications. For use cases requiring higher fidelity, lightweight post-processing (e.g., short minimizations) can be added to improve agreement with the physical model.

The MDZip code is available on github, and each user can apply it to compress their ensembles. Thus, training based on a starting ensemble will produce the trained parameters. Using these parameters, the user can compress the ensemble. Once compressed, the ensembles are easy to share. For anyone to decompress the ensembles, they will need: 1) the trained model parameters, 2) coordinate scalers, 3) molecular topology, and 4) compressed trajectory data. Because the compressed data cannot be interpreted without the trained model, this creates a natural privacy barrier while allowing efficient data distribution. This feature is particularly relevant for collaborations involving proprietary systems or pre-publication datasets where data security is a concern.

The trade-off between local accuracy and full-ensemble accessibility is ultimately application-dependent. In some cases, hybrid strategies may offer the best of both worlds. For example, the MDZip-compressed ensemble can be shared alongside a few restart structures (including velocities) sampled along the trajectory. These high-fidelity structures preserve local geometry and energetic detail and can serve as seeds for regenerating ensembles when physically accurate dynamics are required. For most users, however, the MDZip-compressed ensemble alone may be sufficient for characterizing global conformational properties.

Importantly, the framework integrates seamlessly with existing MD analysis workflows. Once decompressed, the trajectory is restored as a full all-atom ensemble and can be processed with standard tools. With its significant storage and bandwidth savings, MDZip provides a practical, FAIR^39^-compatible solution for molecular simulation data sharing and archival.

## Conclusion

We have introduced *MDZip*, a pair of physics-agnostic convolutional autoencoders that compress molecular-dynamics trajectories by (∼95%) while preserving ensemble fidelity. Trained solely on Cartesian coordinates, both models reproduce global metrics (RMSD, radius of gyration, secondary structure, principal components) and most local features (bond lengths, dihedrals, base-pair parameters) with sub-angstrom accuracy; the skip-connection variant consistently yields fewer outliers and lower errors.

A systematic sweep of latent-space sizes shows diminishing returns for high bottleneck dimensions, providing a practical guide for users to balance storage savings against reconstruction error. Energetic terms remain the main limitation, as small coordinate deviations can give rise to large differences in energetics (especially van der Waals), and a post-processing short minimization strategy helps mitigate this.

MDZip requires only four items for decompression (ensemble-specific trained weights, scalers, topology, and latent trajectory), enabling secure, FAIR-compliant data sharing that fits within resource-constrained repositories such as Zenodo. By lowering the storage barrier without sacrificing atomic detail, our framework offers an immediate, drop-in solution for large-scale MD archiving and analysis.

These results establish a foundation for next-generation trajectory compression methodologies. Future developments integrating physics-aware loss functions or hybrid approaches combining geometric and energetic training objectives hold promise for addressing current limitations while maintaining the computational efficiency and universal applicability demonstrated by our framework.

## Acknolwedgements

This work was supported by the National Science Foundation [CHE-2235785].

## Supporting Information

Supplementary figures 1-21 are provided in SI.

## TOC Graphic

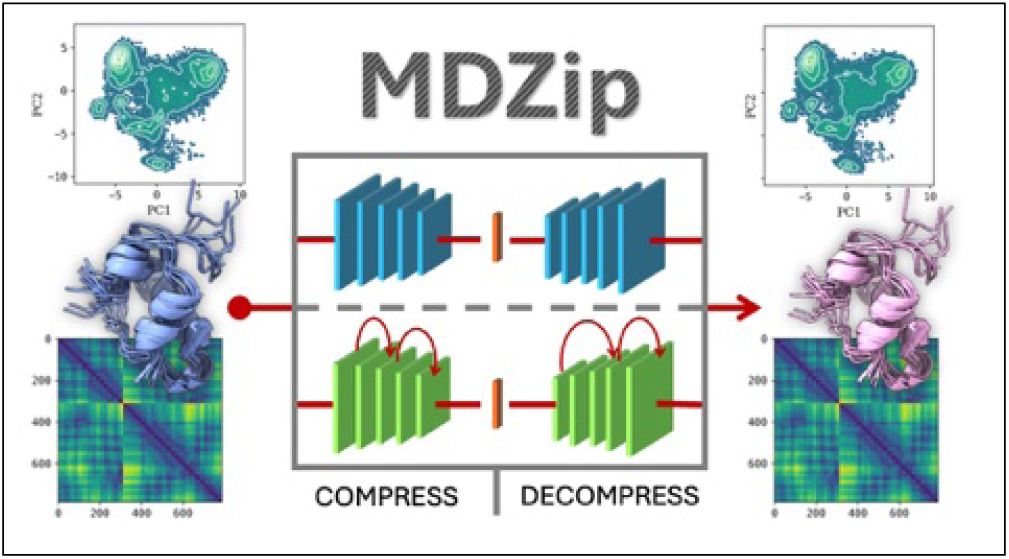

## References

(1) Hospital, A.; Battistini, F.; Soliva, R.; Gelpí, J. L.; Orozco, M. Surviving the deluge of biosimulation data. WIREs Computational Molecular Science 2020, 10, e1449.

(2) Adcock, S. A.; McCammon, J. A. Molecular Dynamics: Survey of Methods for Simulating the Activity of Proteins. Chemical Reviews 2006, 106, 1589–1615, PMID: 16683746.

3. Salo-Ahen, O. M. H. et al. Molecular Dynamics Simulations in Drug Discovery and Pharmaceutical Development. Processes 2021, 9.

(4) Warshel, A. Molecular Dynamics Simulations of Biological Reactions. Accounts of Chemical Research 2002, 35, 385–395, PMID: 12069623.

(5) Tiemann, J. K. S.; Szczuka, M.; Bouarroudj, L.; Oussaren, M.; Garcia, S.; Howard, R. J.; Delemotte, L.; Lindahl, E.; Baaden, M.; Lindorff-Larsen, K.; Chavent, M.; Poulain, P. MDverse, shedding light on the dark matter of molecular dynamics simulations. eLife 2024, 12, RP90061.

(6) Abraham, M. et al. Sharing Data from Molecular Simulations. Journal of Chemical Information and Modeling 2019, 59, 4093–4099.

(7) Amaro, R. et al. The need to implement FAIR principles in biomolecular simulations. arXiv 2024,

(8) Meyer, T.; D’Abramo, M.; Hospital, A.; Rueda, M.; Ferrer-Costa, C.; Perez, A.; Carrillo, O.; Camps, J.; Fenollosa, C.; Repchevsky, D.; Gelpí, J. L.; Orozco, M. MoDEL (Molecular Dynamics Extended Library): a database of atomistic molecular dynamics trajectories. Structure (London, England : 1993) 2010, 18, 1399 – 1409.

(9) Shkurti, A.; Goni, R.; Andrio, P.; Breitmoser, E.; Bethune, I.; Orozco, M.; Laughton, C. A. pyPcazip: A PCA-based toolkit for compression and analysis of molecular simulation data. SoftwareX 2016, 5, 44–50.

(10) Meyer, T.; Ferrer-Costa, C.; Pérez, A.; Rueda, M.; Bidon-Chanal, A.; Luque, F. J.; Laughton, C. A.; Orozco, M. Essential Dynamics: A Tool for Efficient Trajectory Compression and Management. Journal of Chemical Theory and Computation 2006, 2, 251–258, PMID: 26626512.

(11) He, K.; Zhang, X.; Ren, S.; Sun, J. Deep Residual Learning for Image Recognition. CoRR 2015, *abs/1512.03385*.

(12) Peng, Y.; Zhang, L.; Liu, S.; Wu, X.; Zhang, Y.; Wang, X. Dilated Residual Networks with Symmetric Skip Connection for image denoising. Neurocomputing 2019, 345, 67– 76.

(13) B, A.; Kaur, M.; Singh, D.; Roy, S.; Amoon, M. Efficient Skip Connections-Based Residual Network (ESRNet) for Brain Tumor Classification. Diagnostics 2023, 13.

(14) Puliyanda, A.; Devan Padmanathan, A. M.; Mushrif, S. H.; Prasad, V. A 3d convolutional neural network autoencoder for predicting solvent configuration changes in condensed phase biomass reactions. Digital Discovery 2024, 3, 1130–1143.

(15) Kuzminykh, D.; Polykovskiy, D.; Kadurin, A.; Zhebrak, A.; Baskov, I.; Nikolenko, S.; Shayakhmetov, R.; Zhavoronkov, A. 3D Molecular Representations Based on the Wave Transform for Convolutional Neural Networks. Molecular Pharmaceutics 2018, 15, 4378–4385, PMID: 29473756.

(16) Loshchilov, I.; Hutter, F. Decoupled Weight Decay Regularization. 2019; https://arxiv.org/abs/1711.05101.

(17) Eastman, P. et al. OpenMM 4: A Reusable, Extensible, Hardware Independent Library for High Performance Molecular Simulation. Journal of Chemical Theory and Computation 2013, 9, 461–469.

(18) Maier, J. A.; Martinez, C.; Kasavajhala, K.; Wickstrom, L.; Hauser, K. E.; Simmerling, C. ff14SB: Improving the Accuracy of Protein Side Chain and Backbone Parameters from ff99SB. Journal of Chemical Theory and Computation 2015, 11, 3696–3713, PMID: 26574453.

(19) Ivani, I. et al. Parmbsc1: a refined force field for DNA simulations. Nature Methods 2016, 13, 55–58, Epub 2015 Nov 16.

(20) Pérez, ; Marchán, I.; Svozil, D.; Sponer, J.; Cheatham, T. E.; Laughton, C. A.; Orozco, M. Refinement of the AMBER force field for nucleic acids: improving the description of alpha/gamma conformers. Biophysical Journal 2007, 92, 3817–3829, Epub 2007 Mar 9.

(21) McGibbon, R. T.; Beauchamp, K. A.; Harrigan, M. P.; Klein, C.; Swails, J. M.; Hernández, C. X.; Schwantes, C. R.; Wang, L.-P.; Lane, T. J.; Pande, V. S. MDTraj: A Modern Open Library for the Analysis of Molecular Dynamics Trajectories. Biophysical Journal 2015, 109, 1528 – 1532.

(22) Hess, B. Similarities between principal components of protein dynamics and random diffusion. Physical review. E, Statistical physics, plasmas, fluids, and related interdisciplinary topics 2000, 62, 8438 – 8448.

(23) Naritomi, Y.; Fuchigami, S. Slow dynamics in protein fluctuations revealed by time-structure based independent component analysis: The case of domain motions. The Journal of Chemical Physics 2011, 134, 065101.

(24) Pedregosa, F. et al. Scikit-learn: Machine Learning in Python. Journal of Machine Learning Research 2011, 12, 2825–2830.

(25) Hoffmann, M.; Scherer, M. K.; Hempel, T.; Mardt, A.; de Silva, B.; Husic, B. E.; Klus, S.; Wu, H.; Kutz, J. N.; Brunton, S.; Noé, F. Deeptime: a Python library for machine learning dynamical models from time series data. Machine Learning: Science and Technology 2021,

(26) Roe, D. R.; III, T. E. C. PTRAJ and CPPTRAJ: Software for Processing and Analysis of Molecular Dynamics Trajectory Data. Journal of chemical theory and computation 2013, 9, 3084 – 3095.

(27) Case, D. et al. AMBER 2020. 2020.

(28) Darden, T.; York, D.; Pedersen, L. Particle mesh Ewald: An Nlog(N) method for Ewald sums in large systems. The Journal of chemical physics 1993, 98, 10089 – 10092.

(29) Lu, X.; Olson, W. K. 3DNA: a software package for the analysis, rebuilding and visualization of three-dimensional nucleic acid structures. Nucleic Acids Research 2003, 31, 5108–5121.

(30) Olson, W. K. et al. A standard reference frame for the description of nucleic acid base-pair geometry11Edited by P. E. Wright22This is a document of the Nomenclature Committee of IUBMB (NC-IUBMB)/IUPAC-IUBMB Joint Commission on Biochemical Nomenclature (JCBN), whose members are R. Cammack (chairman), A. Bairoch, H.M. Berman, S. Boyce, C.R. Cantor, K. Elliott, D. Horton, M. Kanehisa, A. Kotyk, G.P. Moss, N. Sharon and K.F. Tipton. Journal of Molecular Biology 2001, 313, 229–237.

(31) Olson, W. K.; Gorin, A. A.; Lu, X. J.; Hock, L. M.; Zhurkin, V. B. DNA sequence-dependent deformability deduced from protein-DNA crystal complexes. Proceedings of the National Academy of Sciences 1998, 95, 11163 – 11168.

(32) European Organization For Nuclear Research; OpenAIRE Zenodo. 2013; https://www.zenodo.org/.

(33) Acun, B.; Hardy, D. J.; Kale, L. V.; Li, K.; Phillips, J. C.; Stone, J. E. Scalable molecular dynamics with NAMD on the Summit system. IBM Journal of Research and Development 2018, 62, 4:1–4:9.

(34) Chipot, C. Recent Advances in Simulation Software and Force Fields: Their Importance in Theoretical and Computational Chemistry and Biophysics. The Journal of Physical Chemistry B 2024, 128, 12023–12026, PMID: 39663898.

(35) Jung, J.; Nishima, W.; Daniels, M.; Bascom, G.; Kobayashi, C.; Adedoyin, A.; Wall, M.; Lappala, A.; Phillips, D.; Fischer, W.; Tung, C.-S.; Schlick, T.; Sugita, Y.; Sanbonmatsu, K. Y. Scaling molecular dynamics beyond 100,000 processor cores for large-scale biophysical simulations. Journal of Computational Chemistry 2019, 40, 1919–1930, Epub 2019 Apr 17.

(36) Miyamoto, S.; Kollman, P. A. SETTLE: an analytical version of the SHAKE and RATTLE algorithm for rigid water models. Journal of Computational Chemistry 1992, 13, 952-962.

(37) Elber, R.; Ruymgaart, A. P.; Hess, B. SHAKE parallelization. The European Physical Journal Special Topics 2011, 200, 211–223.

(38) Kabsch, W.; Sander, C. Dictionary of protein secondary structure: Pattern recognition of hydrogen-bonded and geometrical features. Biopolymers 1983, 22, 2577–2637.

(39) Wilkinson, M. D. et al. The FAIR Guiding Principles for scientific data management and stewardship. Scientific Data 2016, 3, 160018, Erratum in: Sci Data. 2019 Mar 19;6(1):6. doi: 10.1038/s41597-019-0009-6.

