## Supplementary material for "MDZip: Neural Compression of Molecular Dynamics Trajectories for Scalable Storage and Ensemble Reconstruction": SI figures

### SI: MDZip: Efficient data compression for MD data sharing

Namindu De Silva[1], Alberto Perez[1,*]
[1]Department of Chemistry, University of Florida, Gainesville, FL 32611, USA
*

July 31, 2025

#### 1 Supplementary figures

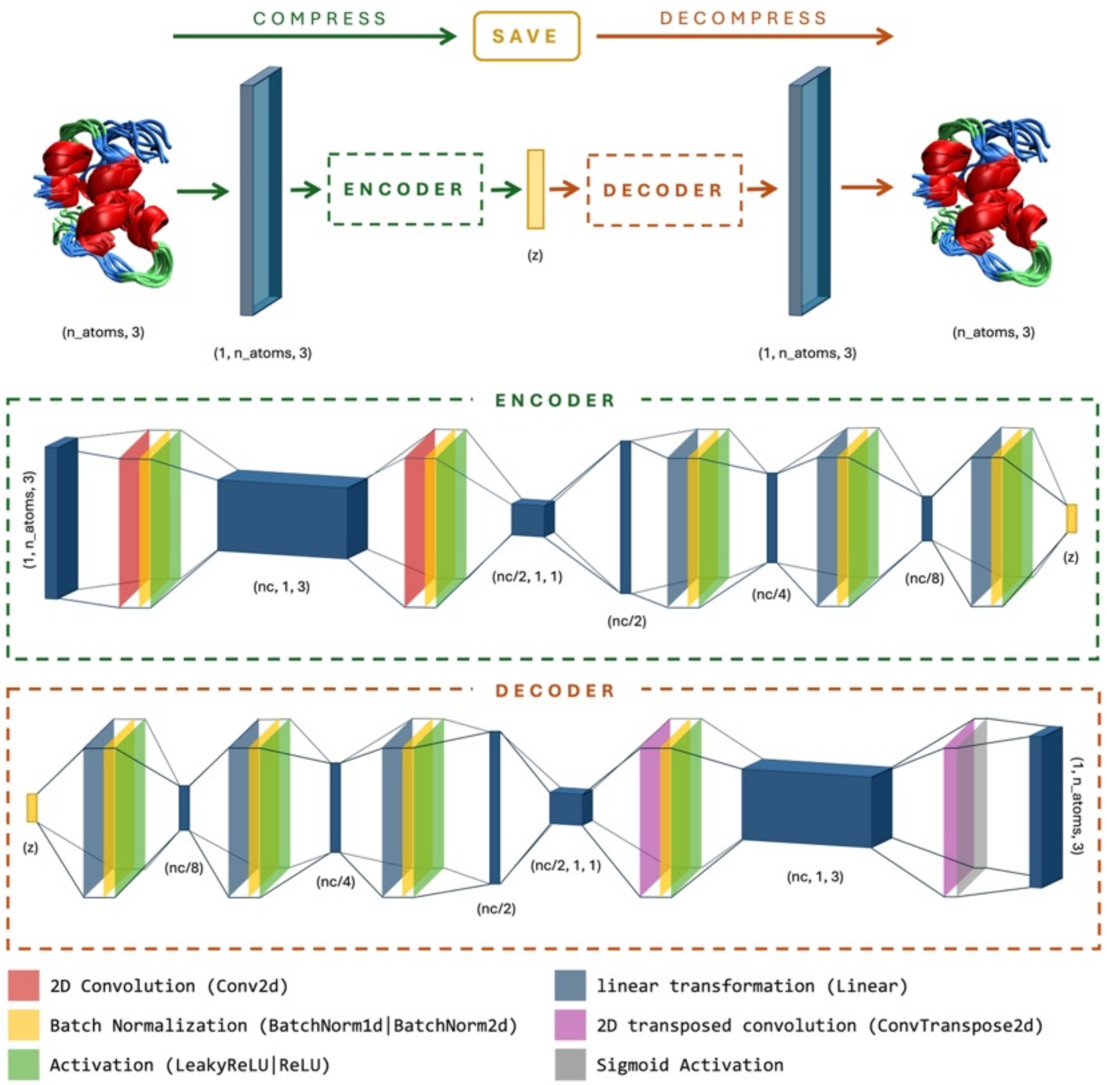


SI Figure 1: Basic architecture of the autoencoder used to compress molecular dynamics (MD) trajectory data. Input, output and intermediate tensor dimensions are indicated in parentheses. Here, $n\_atoms$ denotes the number of atoms in the trajectory, and the channel count $nc$ is set to 1024 if $3 * n\_atoms < 1024$, to 2048 if $1024 \leq 3 * n_atoms < 2048$, and to 4096 (default) or a user-specified value otherwise.

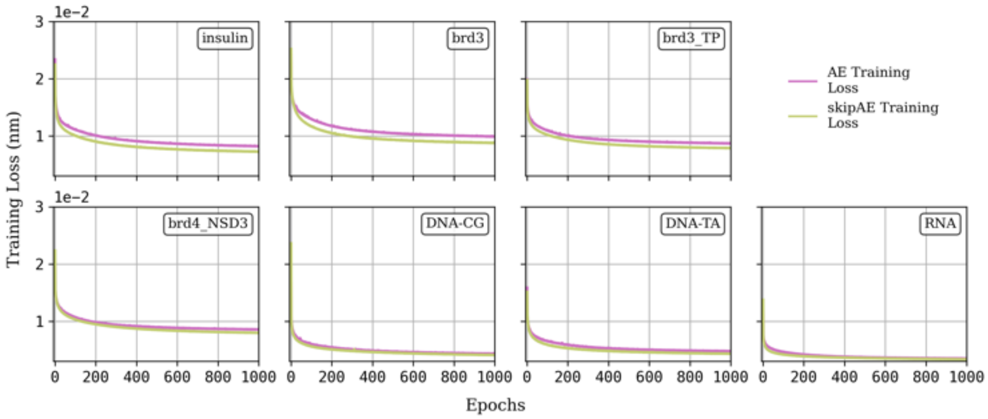


SI Figure 2: Training loss curves for conventional autoencoder (AE, red) and skip-connection enhanced autoencoder (skipAE, green) across all molecular systems. Both architectures demonstrate smooth convergence within 1000 epochs, reaching stable performance plateaus without oscillatory behavior.

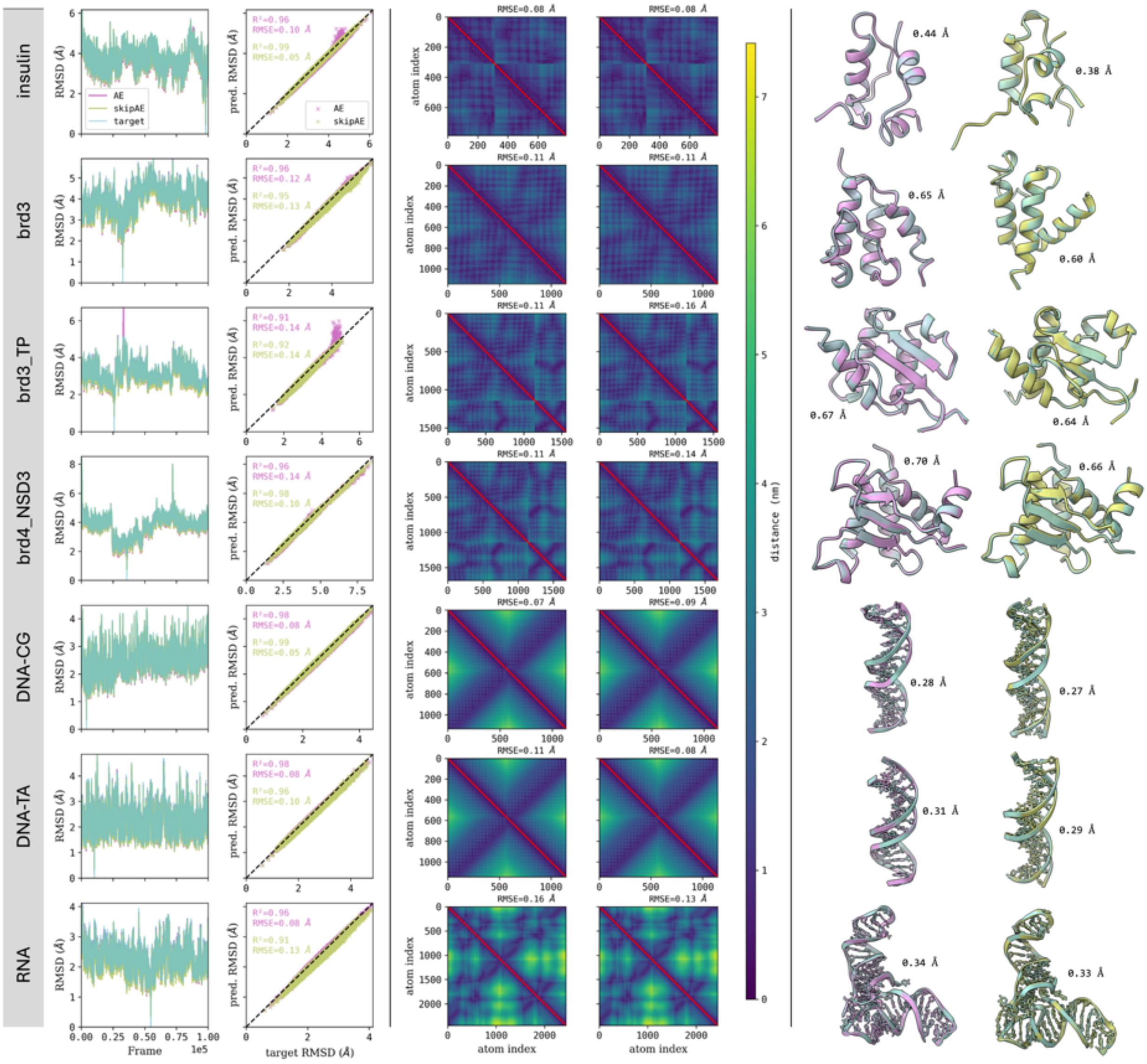

SI Figure 3: Evaluation of autoencoder reconstruction performance across molecular systems. Each row corresponds to a distinct molecular system. (Column 1) RMSD fluctuations for target (cyan), AE (magenta), and skipAE (green) trajectories. (Column 2) Target versus predicted RMSD scatter plots with RMSE and $R^2$ values in corresponding colors. (Columns 3-4) Mean distance matrices showing target (upper triangle) and predicted (lower triangle) distances for AE and skipAE models, respectively, with RMSE values at top right. (Columns 5-6) Best reconstructed structures (lowest $\Delta RMSD$) for target (cyan), AE (magenta), and skipAE (green), with $\Delta RMSD$ values displayed for quantitative comparison.

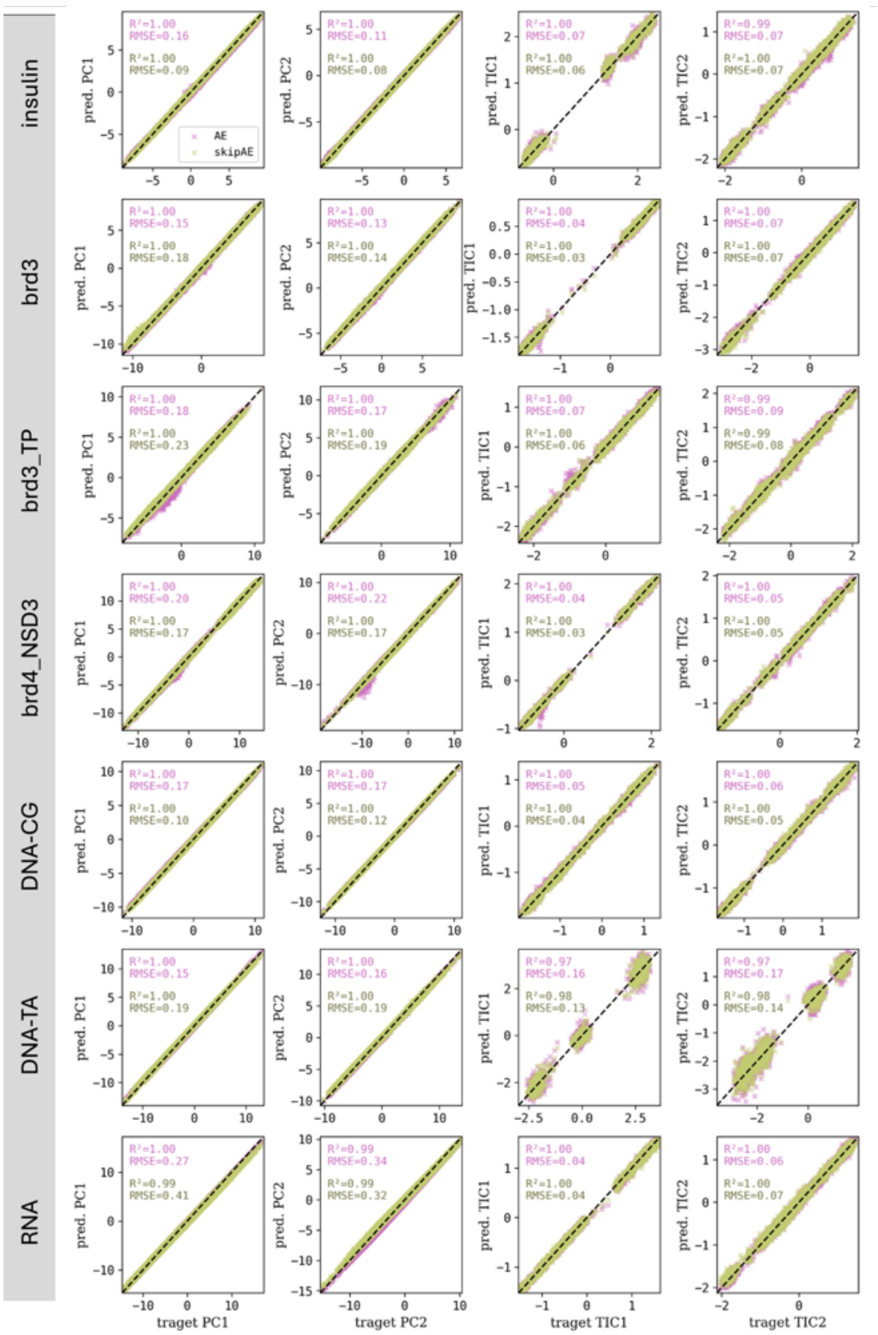


SI Figure 4: Correlation analysis of principal and independent component projections between target and reconstructed trajectories. Each row represents a distinct molecular system. Columns 1-4 show predicted versus target values for PC1, PC2, TIC1, and TIC2, respectively, with conventional autoencoder (AE, magenta) and skip-connection enhanced autoencoder (skipAE, green) predictions. $R^2$ and RMSE values are displayed in the top left corner in corresponding model colors, demonstrating near-perfect correlation between predicted and target dynamical projections.

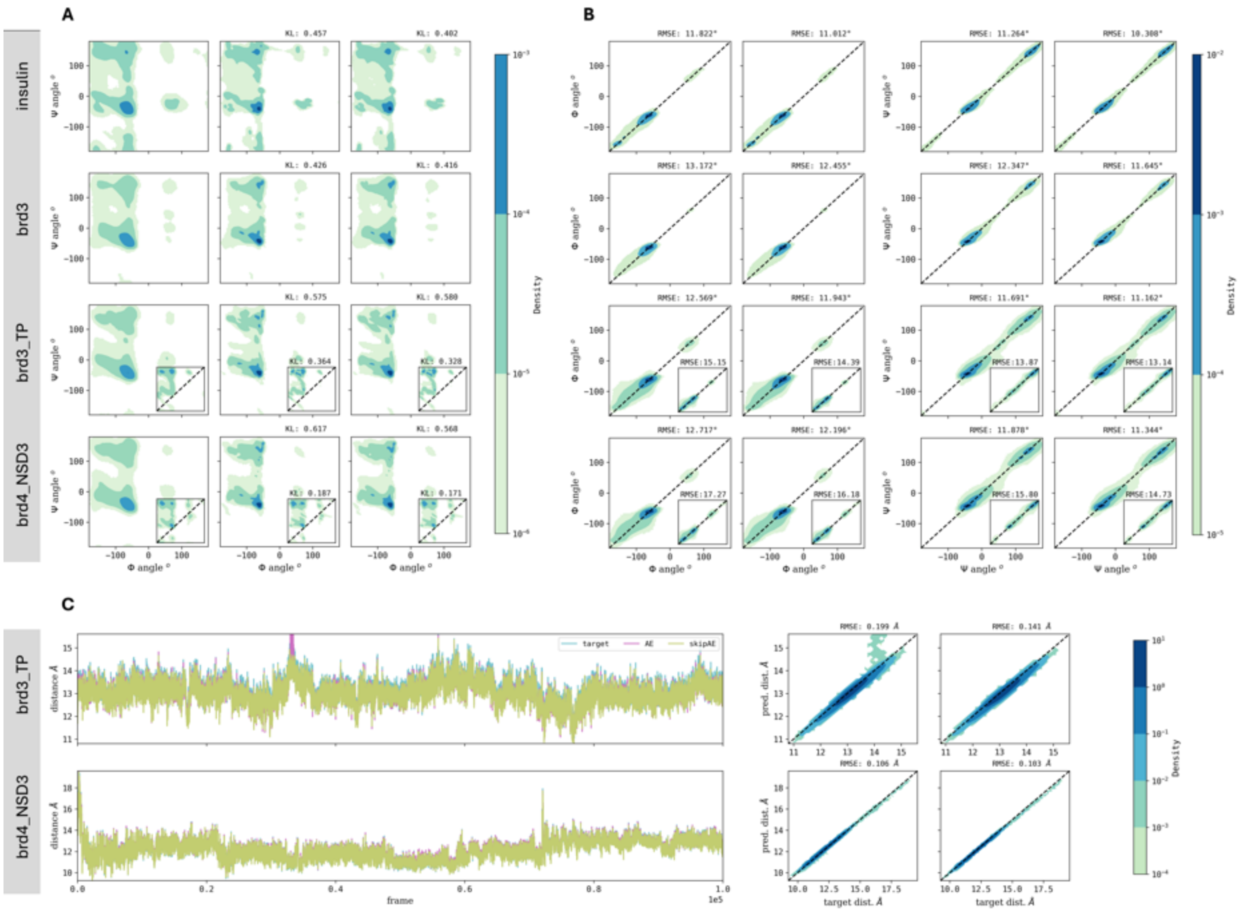


SI Figure 5: Ramachandran plot analysis of backbone dihedral angle preservation across protein systems. Each row represents a distinct protein system as labeled. **(A)** Ramachandran plots showing the distribution of Ψ versus Φ angles for target structures (column 1), AE predictions (column 2), and skipAE predictions (column 3). For protein-peptide complexes, main plots display protein distributions while small inset plots show peptide distributions. Kullback-Leibler (KL) divergence values, calculated with respect to the target distribution, are displayed at the top right of each plot. **(B)** Correlation analysis of predicted versus target Φ angles for AE (column 1) and skipAE (column 2) models. **(C)** Correlation analysis of predicted versus target Ψ angles for AE (column 1) and skipAE (column 2) models. For panels B and C, protein-peptide complexes display peptide angle correlations in small inset plots within the main protein plots. RMSE values are shown at the top right of each correlation plot. **(C)** Center-of-mass distance analysis for protein-peptide complexes. Each row represents a distinct complex. (Column 1) Distance fluctuations throughout the trajectory for target (cyan), AE (magenta), and skipAE (green). (Columns 2-3) Correlation plots of predicted versus target COM distances for AE and skipAE models, respectively. RMSE values are shown at the top right of each plot.

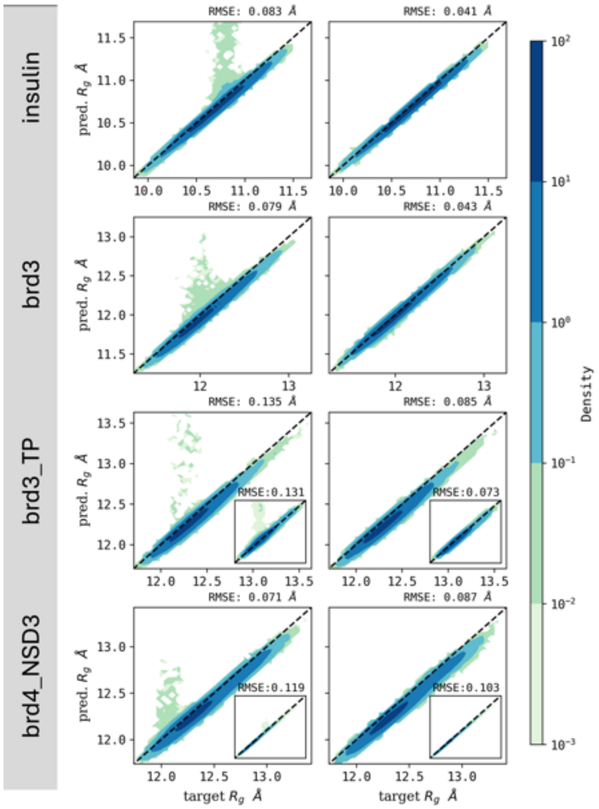


SI Figure 6: Radius of gyration correlation analysis showing predicted versus target values across protein systems. Each row represents a distinct protein system as labeled. **(Column 1)** AE predictions versus target radius of gyration values. **(Column 2)** SkipAE predictions versus target radius of gyration values. For protein-peptide complexes, peptide radius of gyration correlations are displayed in small inset plots within the main protein plots. RMSE values are shown for quantitative assessment of reconstruction accuracy.

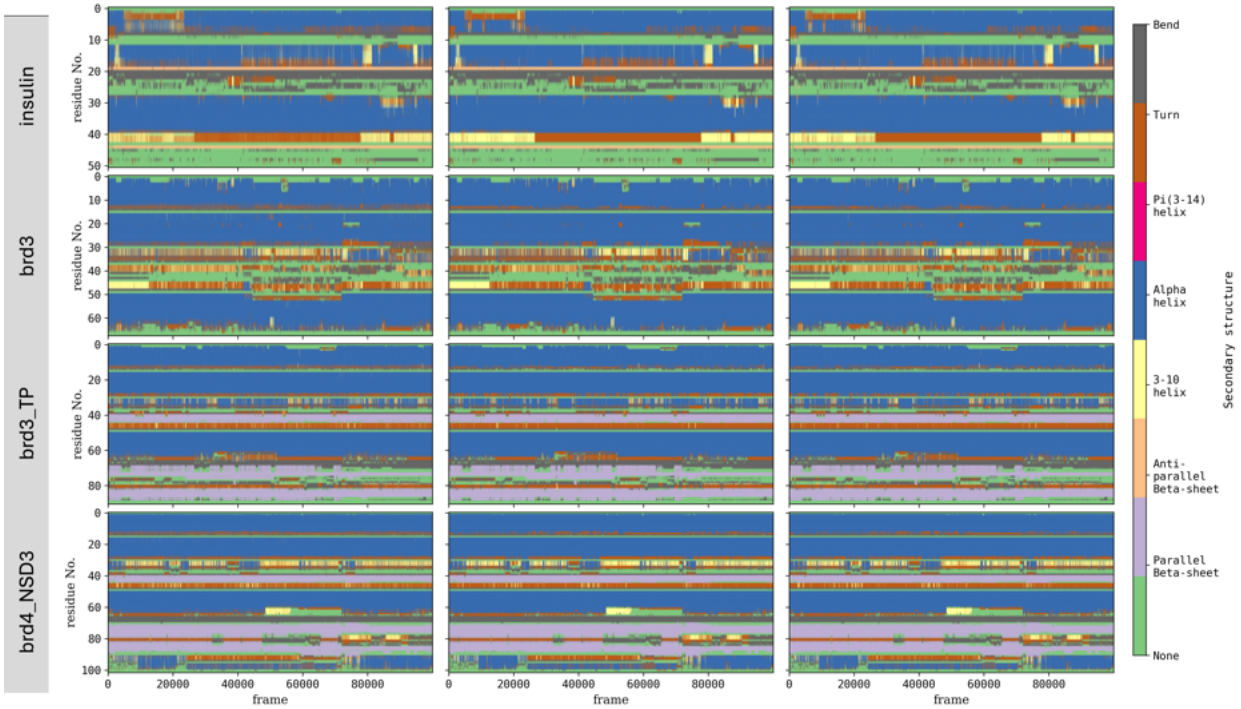


SI Figure 7: Secondary structure preservation analysis using DSSP (Dictionary of Secondary Structure of Proteins) assignments across protein systems. Each row represents a distinct protein system as labeled. **(Column 1)** Target secondary structure evolution throughout the simulation trajectory. **(Column 2)** Conventional autoencoder (AE) predicted secondary structure assignments. **(Column 3)** Skip-connection enhanced autoencoder (skipAE) predicted secondary structure assignments. Color coding represents different secondary structure elements (bend (gray), turn (brown), pi(3-14) helix (pink), $\alpha$-helix (blue), 3-10 helix (yellow), anti-parallel $\beta$-sheets (orange), parallel $\beta$-sheets (purple)) as defined by DSSP criteria, demonstrating preservation of characteristic structural motifs in reconstructed trajectories.

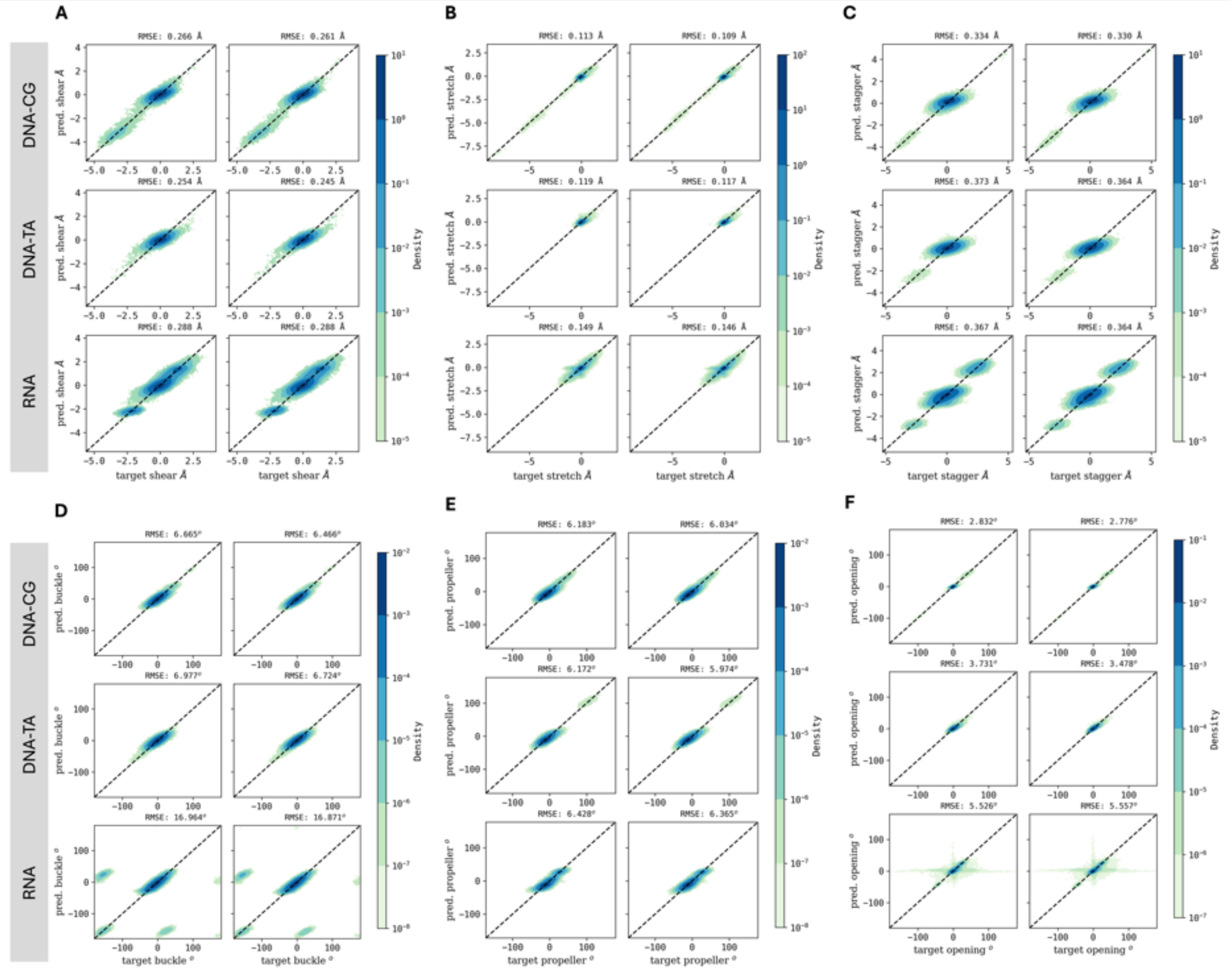


SI Figure 8: Base-pair parameter reconstruction analysis for nucleic acid systems. Each panel represents a distinct base-pair parameter: **(A)** shear, **(B)** stretch, **(C)** stagger, **(D)** buckle, **(E)** propeller, and **(F)** opening. Within each panel, rows correspond to DNA-CG (top), DNA-TA (middle), and RNA (bottom) systems. Columns show correlation plots of predicted versus target values for conventional autoencoder (AE, left) and skip-connection enhanced autoencoder (skipAE, right) models. RMSE values are displayed at the top right of each correlation plot, demonstrating reconstruction accuracy for fundamental nucleic acid structural parameters.

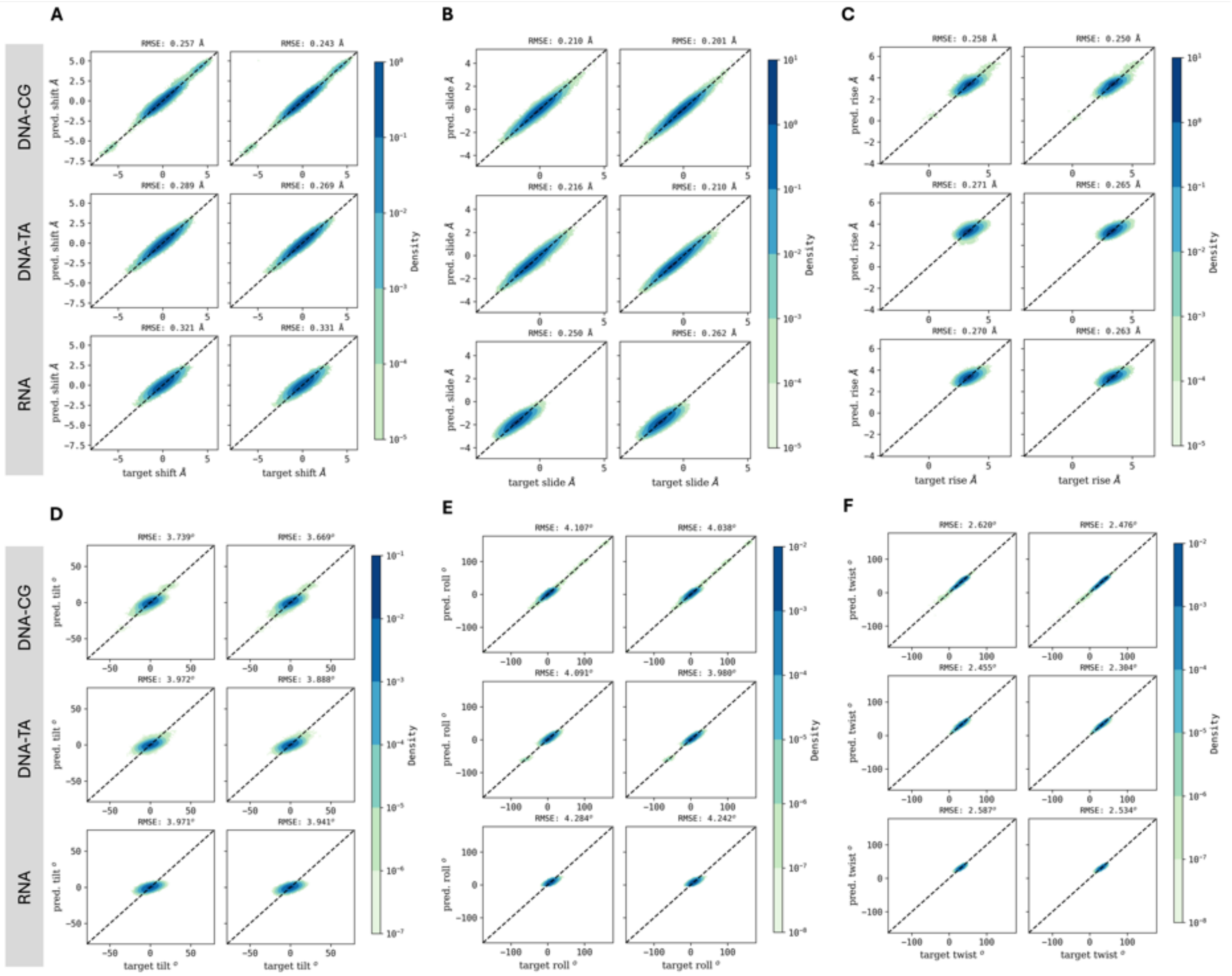


SI Figure 9: Base-pair step parameter reconstruction analysis for nucleic acid systems. Each panel represents a distinct base-pair step parameter: **(A)** shift, **(B)** slide, **(C)** rise, **(D)** tilt, **(D)** roll, and **(F)** twist. Within each panel, rows correspond to DNA-CG (top), DNA-TA (middle), and RNA (bottom) systems. Columns show correlation plots of predicted versus target values for conventional autoencoder (AE, left) and skip-connection enhanced autoencoder (skipAE, right) models. RMSE values are displayed at the top right of each correlation plot, demonstrating reconstruction accuracy for nucleic acid step geometry parameters.

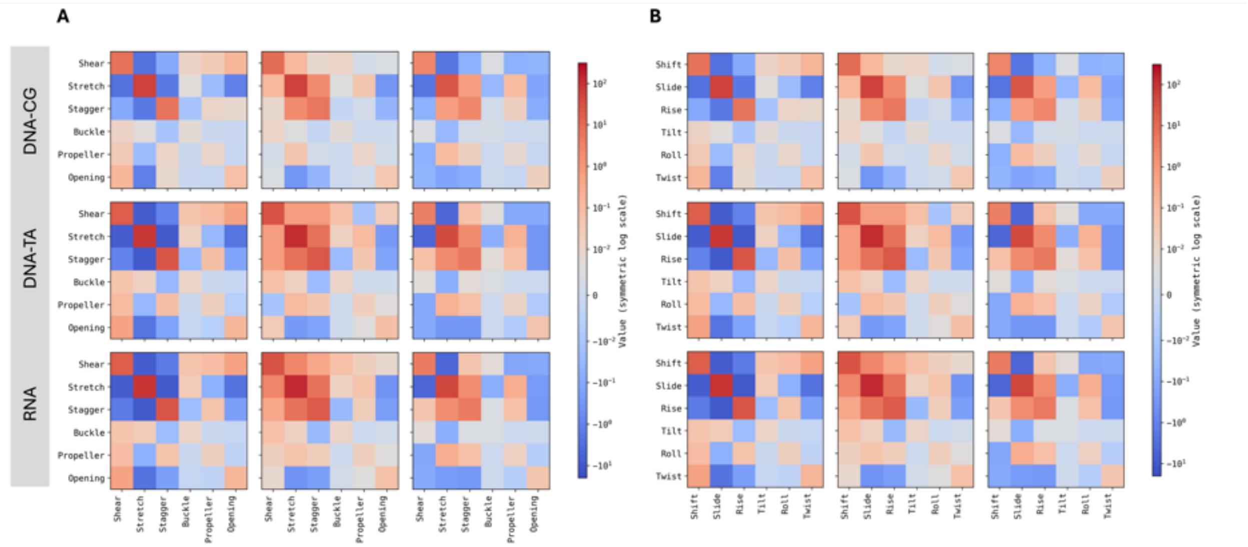


SI Figure 10: Force constant matrices derived from inverted covariance matrices of nucleic acid structural parameters. **(A)** Base-pair parameter force constants and **(B)** base-pair step parameter force constants. Each row represents a distinct molecular system as labeled. Columns display target force constants (left), conventional autoencoder predictions (AE, middle), and skip-connection enhanced autoencoder predictions (skipAE, right). Higher values indicate lower deviation from mean structural values, analogous to stronger force constants.

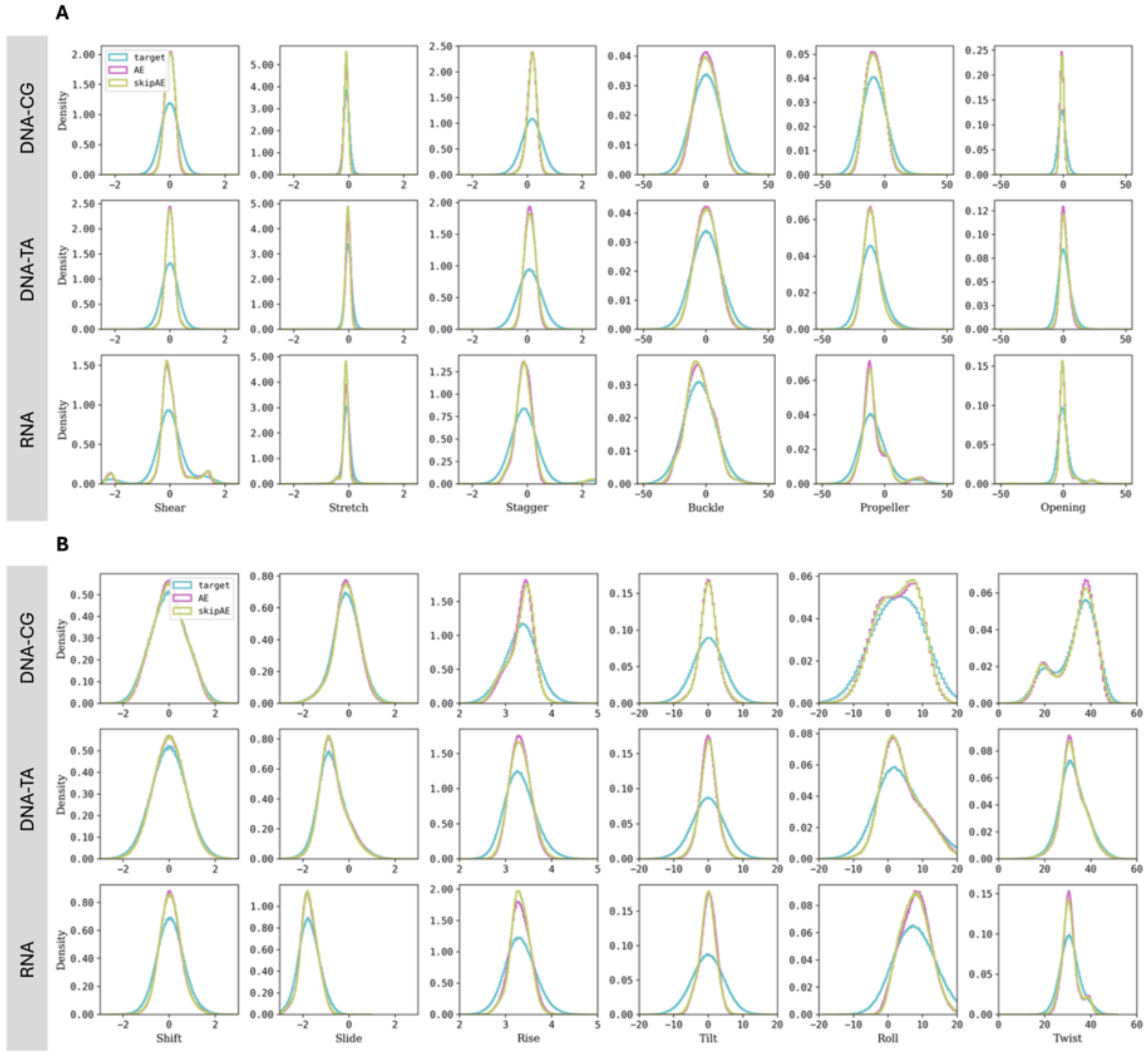


SI Figure 11: Distribution analysis of nucleic acid structural parameters. **(A)** Base-pair parameter distributions and **(B)** base-pair step parameter distributions. Each row represents a distinct molecular system as labeled. Histograms show target distributions (cyan), conventional autoencoder predictions (AE, magenta), and skip-connection enhanced autoencoder predictions (skipAE, green). Kullback-Leibler divergence values quantifying deviation from target distributions are displayed at the top right of each plot in corresponding model colors (AE: magenta, skipAE: green).

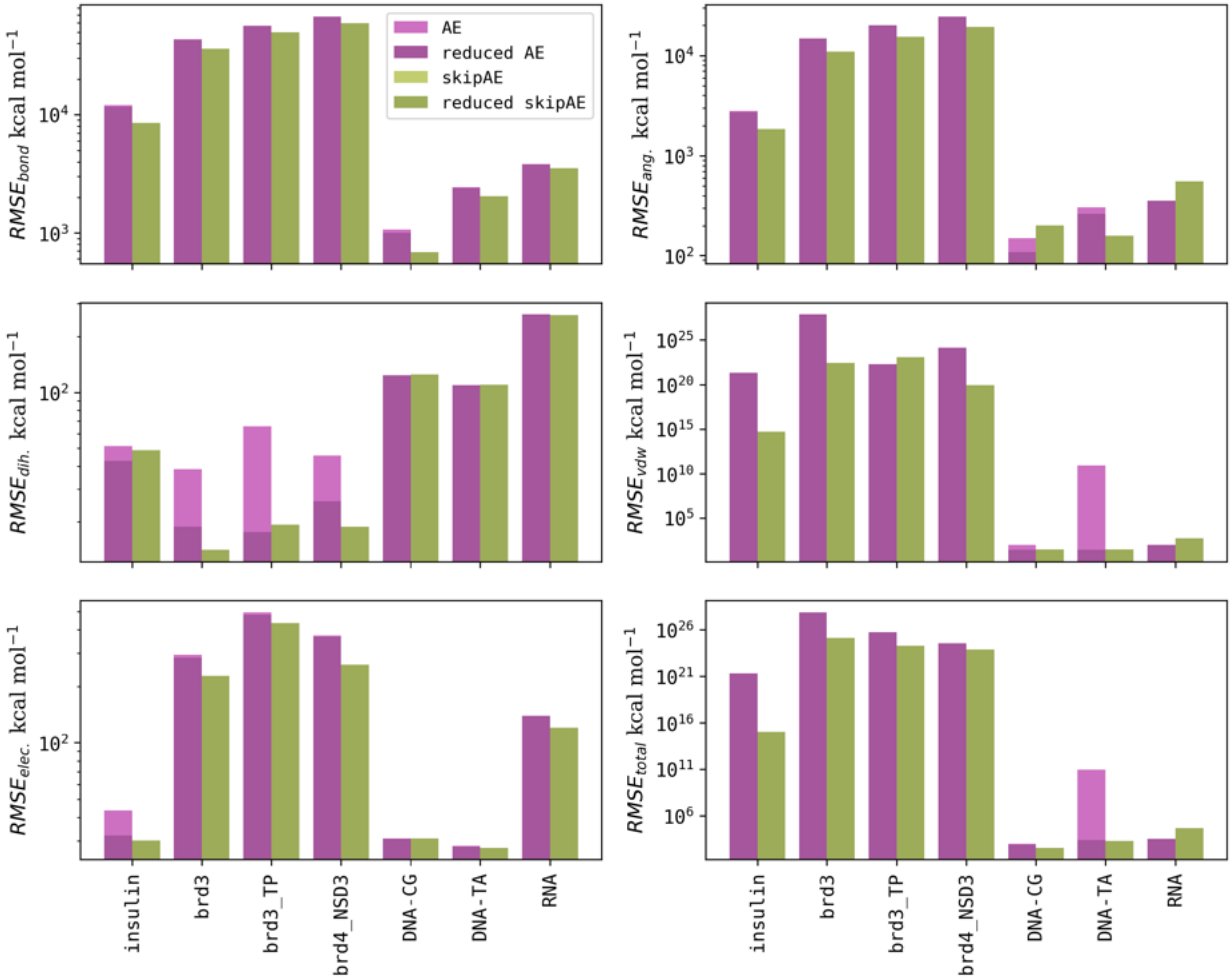


SI Figure 12: Energy component reconstruction analysis comparing all predicted structures versus high-quality filtered structures. (Row 1, Columns 1-2) Bond energy and angle energy RMSE values. (Row 2, Columns 1-2) Dihedral energy and van der Waals energy RMSE values. (Row 3, Columns 1-2) Electrostatic energy and total energy RMSE values. Color coding represents: conventional autoencoder predictions for all structures (magenta) and filtered structures with $\Delta RMSD < 1\AA$ for proteins and $< 0.5\AA$ for DNA/RNA systems (dark magenta); skip-connection enhanced autoencoder predictions for all structures (green) and filtered high-quality structures (dark green). Results demonstrate substantial improvement in energy reconstruction when analysis is restricted to geometrically well-reconstructed conformations.

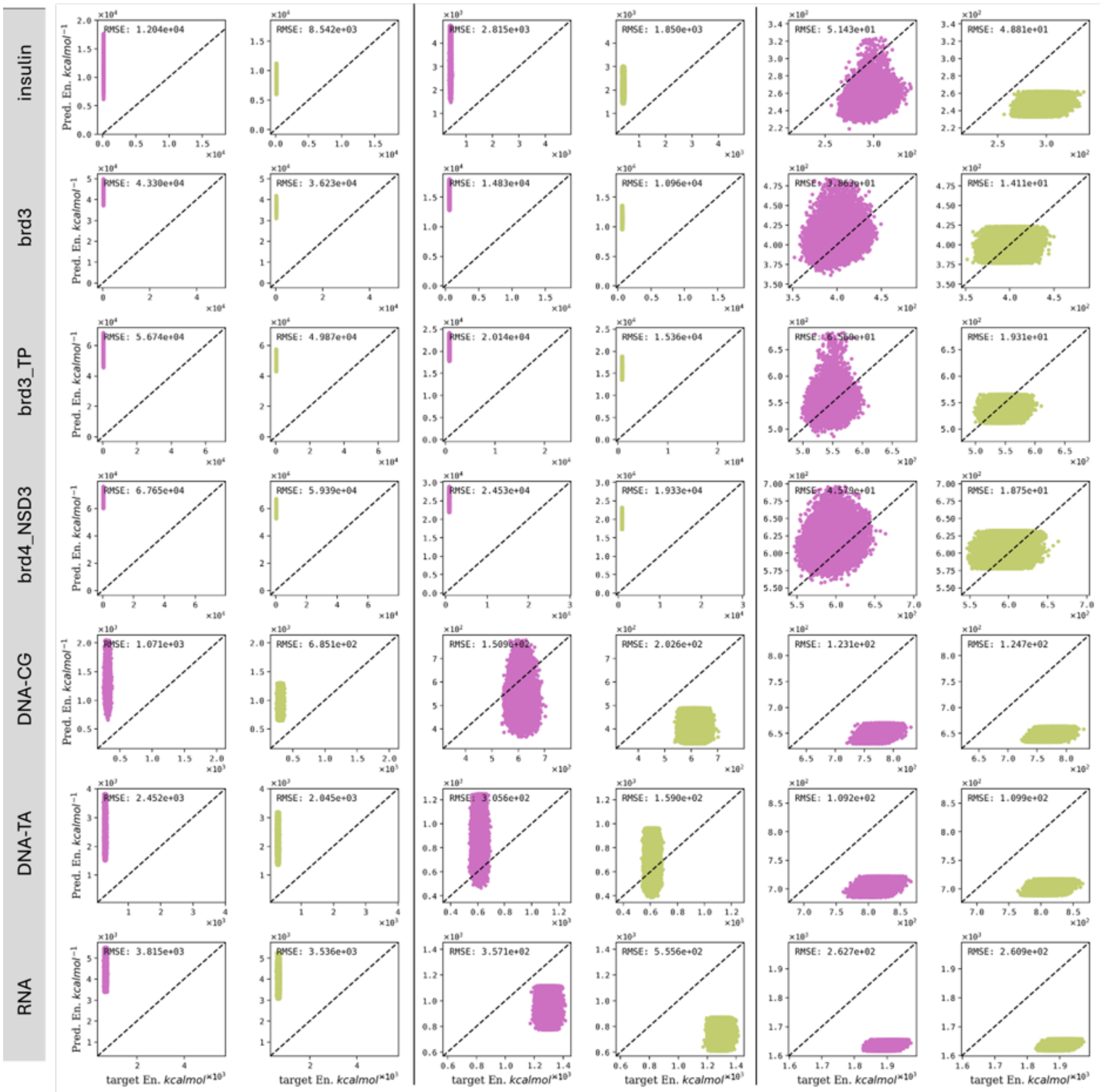


SI Figure 13: Energy component reconstruction analysis across molecular systems. Each row represents a distinct molecular system as labeled. (Columns 1-2) Bond energy correlation plots showing predicted versus target values for conventional autoencoder (AE, column 1) and skip-connection enhanced autoencoder (skipAE, column 2). (Columns 3-4) Angle energy correlation plots for AE (column 3) and skipAE (column 4). (Columns 5-6) Dihedral energy correlation plots for AE (column 5) and skipAE (column 6). RMSE values are displayed at the top right of each correlation plot, demonstrating the challenges in reconstructing energetic properties from purely geometric learning approaches.

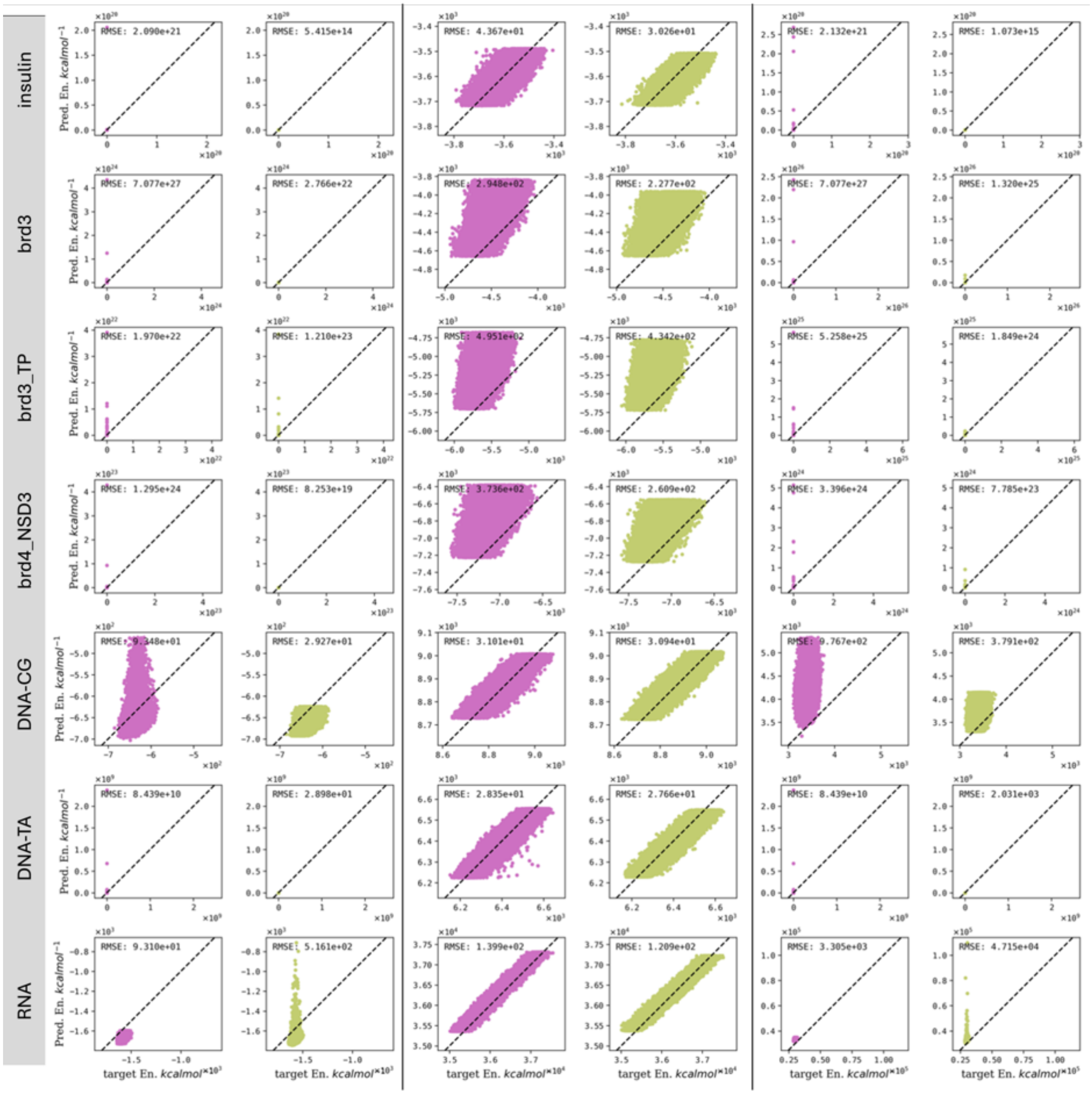


SI Figure 14: Non-bonded and total energy reconstruction analysis across molecular systems. Each row represents a distinct molecular system as labeled. (Columns 1-2) Van der Waals energy correlation plots showing predicted versus target values for conventional autoencoder (AE, column 1) and skip-connection enhanced autoencoder (skipAE, column 2). (Columns 3-4) Electrostatic energy correlation plots for AE (column 3) and skipAE (column 4). (Columns 5-6) Total energy correlation plots for AE (column 5) and skipAE (column 6). RMSE values are displayed at the top right of each correlation plot, illustrating the systematic challenges in energy reconstruction from coordinate-based learning methods.

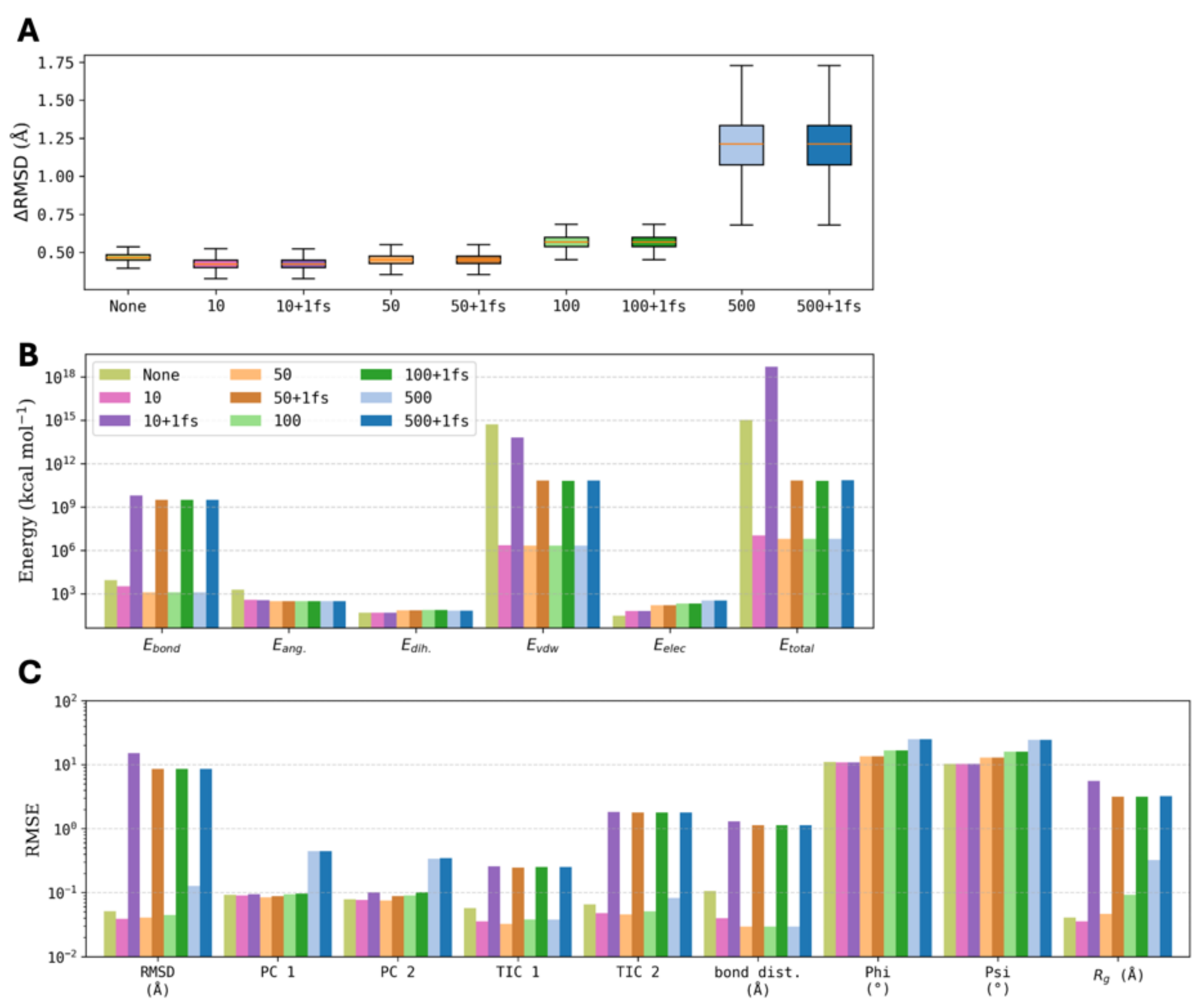


SI Figure 15: Impact of energy minimization protocols on reconstruction accuracy for insulin trajectories. **(A)** Structural deviation ($\Delta RMSD$) from target conformations across different minimization step protocols. **(B)** Energy component reconstruction accuracy showing RMSE values for bonded, non-bonded, and total energies. **(C)** RMSE values for various structural properties including bond lengths, dihedral angles, and radius of gyration. Bar charts demonstrate the trade-off between energy optimization and structural preservation across minimization protocols ranging from 10 to 500 steps.(see SI Fig 16-18 for detailed property descriptions)

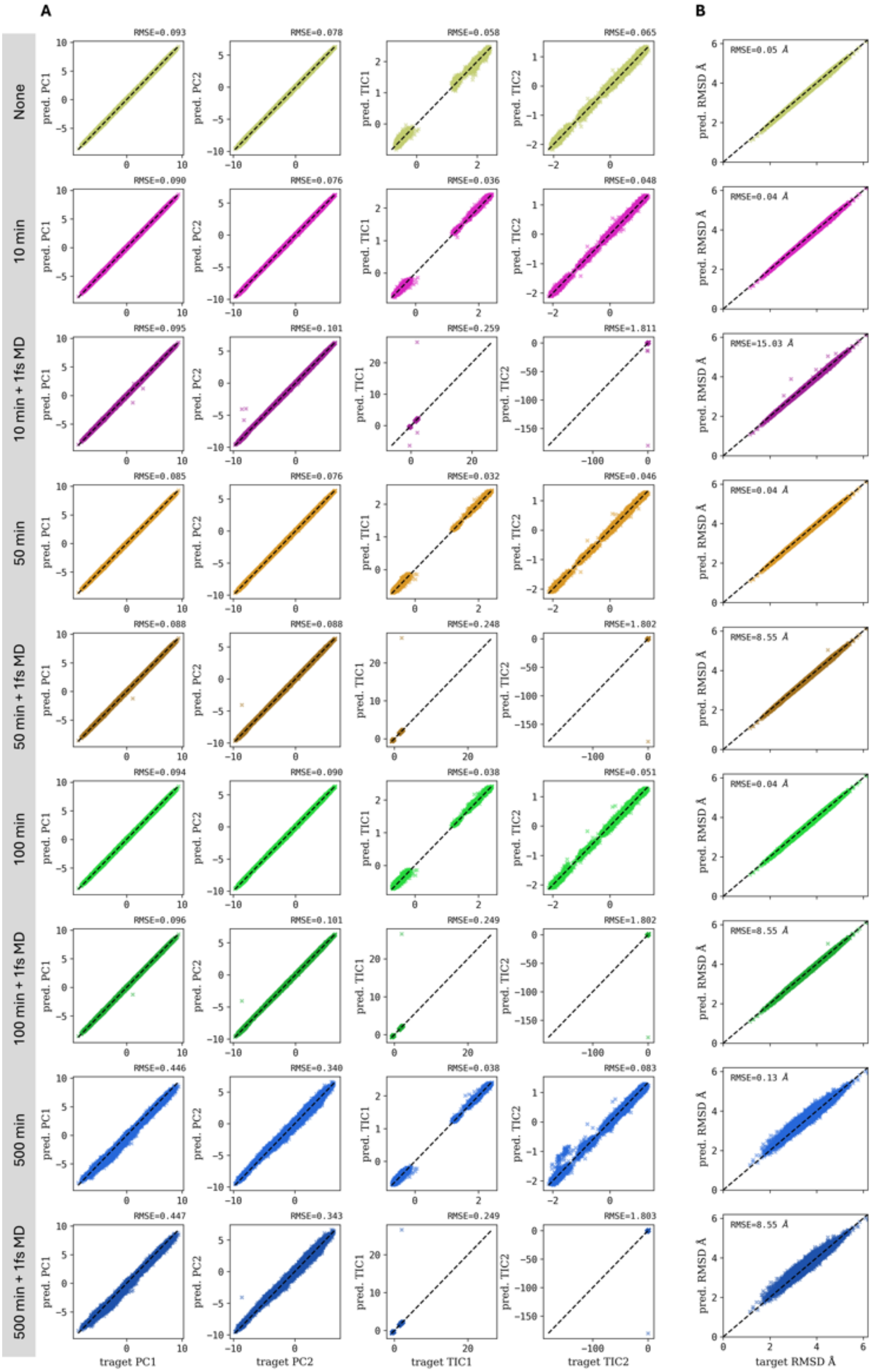


SI Figure 16: Global property preservation analysis following energy minimization protocols. (A) Principal component and time-lagged independent component correlation analysis. Each row represents a distinct minimization protocol. Columns show correlation plots of predicted versus target values for first principal component (PC1, column 1), second principal component (PC2, column 2), first time-lagged independent component (TIC1, column 3), and second time-lagged independent component (TIC2, column 4). (B) Overall structural correlation showing predicted versus target RMSD values.

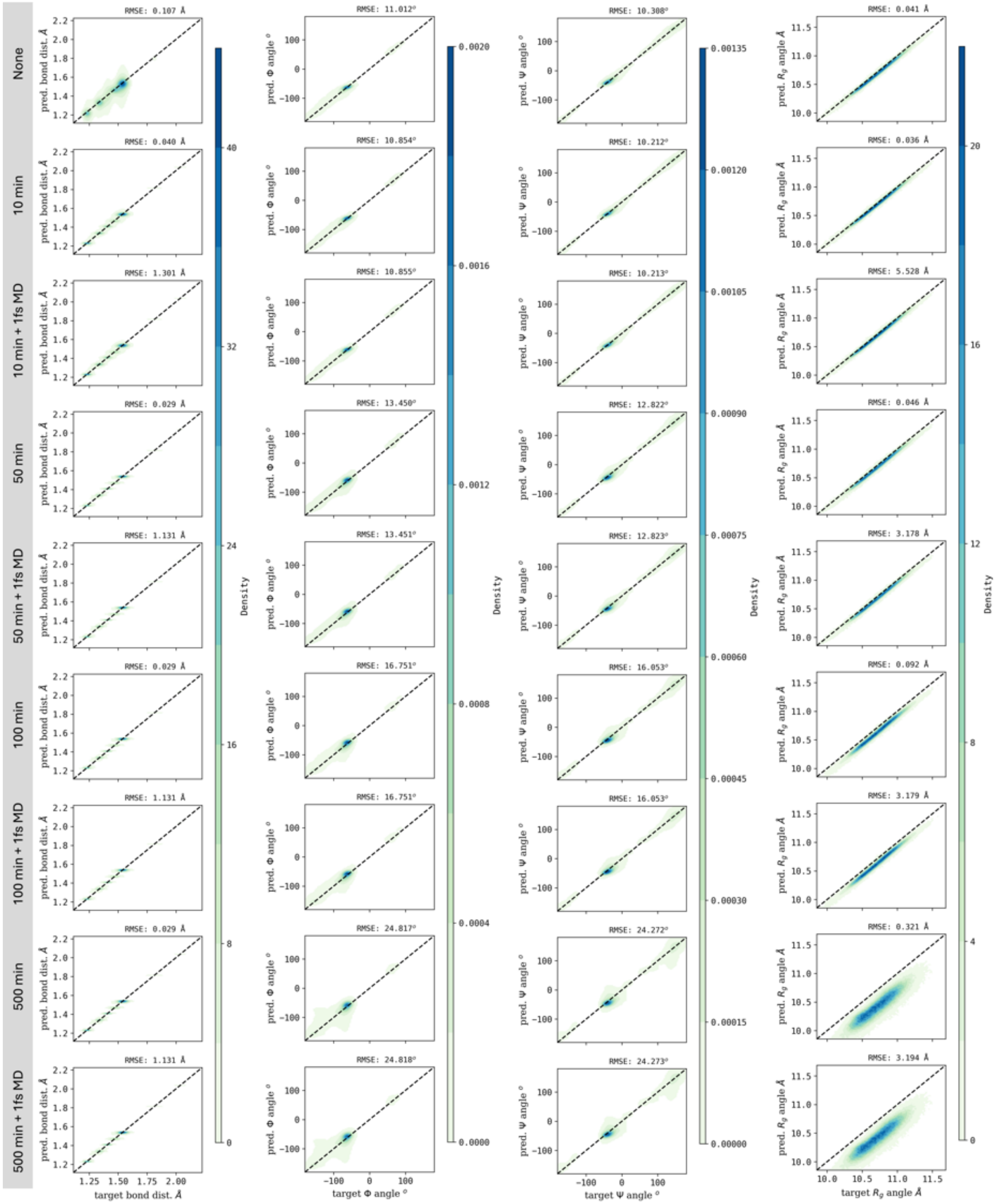


SI Figure 17: Structural property preservation analysis following energy minimization protocols. Each row represents a distinct minimization protocol. Columns show correlation plots of predicted versus target values for bond distances (column 1), Φ dihedral angles (column 2), Ψ dihedral angles (column 3), and radius of gyration (column 4). RMSE values are displayed at the top right of each correlation plot, demonstrating the impact of minimization steps on local and global structural accuracy.

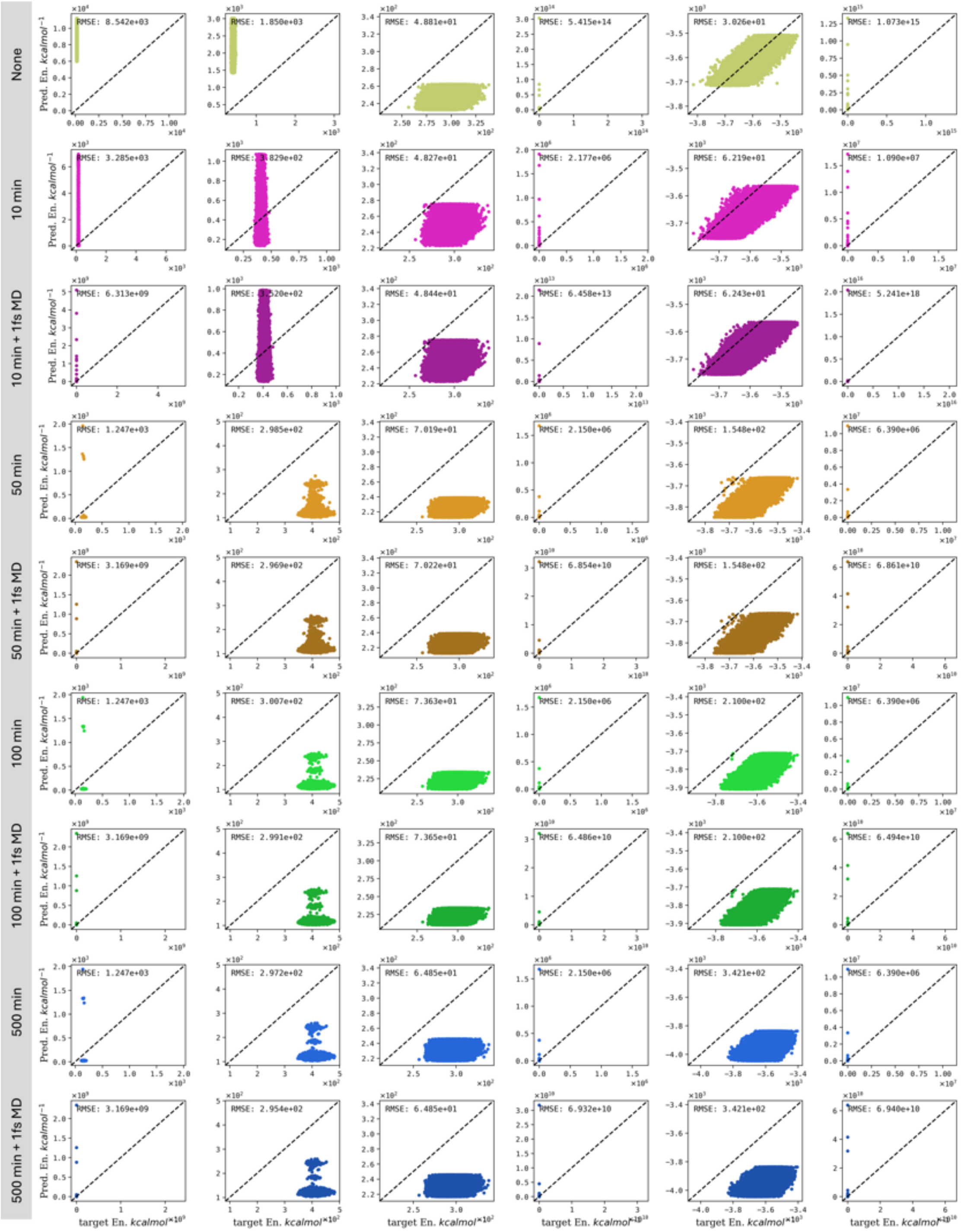


SI Figure 18: Energy component reconstruction analysis following energy minimization protocols. Each row represents a distinct minimization protocol. Columns show correlation plots of predicted versus target values for bond energy (column 1), angle energy (column 2), dihedral energy (column 3), van der Waals energy (column 4), electrostatic energy (column 5), and total energy (column 6). RMSE values are displayed at the top right of each correlation plot, demonstrating the effectiveness of minimization in improving energy reconstruction accuracy across different force field components.

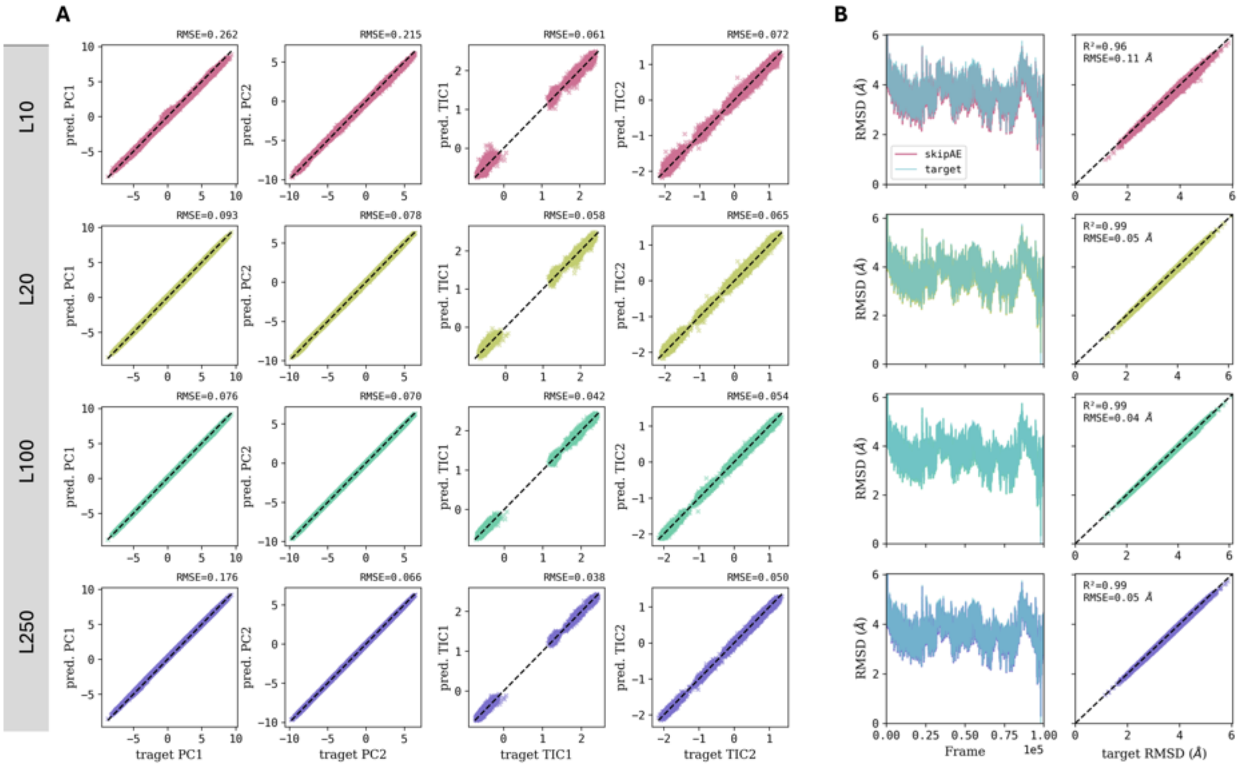


SI Figure 19: Impact of bottleneck dimensionality on dynamical property reconstruction. (A) Principal component and time-lagged independent component correlation analysis across different latent dimensions. Each row represents a distinct bottleneck size as labeled (L10: 10 dimensions, L20: 20 dimensions, L100: 100 dimensions, L250: 250 dimensions). Columns show correlation plots of predicted versus target values for first principal component (PC1, column 1), second principal component (PC2, column 2), first time-lagged independent component (TIC1, column 3), and second time-lagged independent component (TIC2, column 4). (B) RMSD analysis showing fluctuations relative to average structure throughout the simulation trajectory (column 1) and correlation of predicted versus target RMSD values (column 2). Results demonstrate improved dynamical feature preservation with increasing bottleneck dimensions.

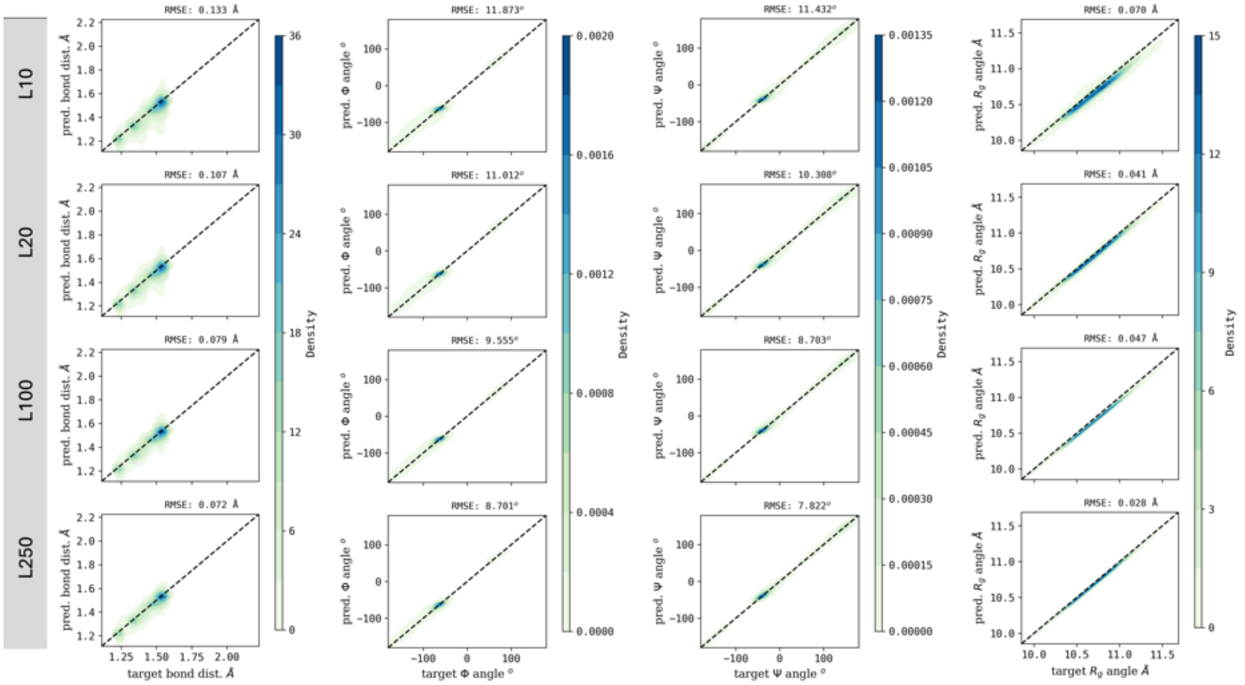


SI Figure 20: Structural property preservation analysis across different bottleneck dimensions. Each row represents a distinct latent dimension as labeled (L10: 10 dimensions, L20: 20 dimensions, L100: 100 dimensions, L250: 250 dimensions). Columns show correlation plots of predicted versus target values for bond distances (column 1), Φ dihedral angles (column 2), Ψ dihedral angles (column 3), and radius of gyration (column 4). RMSE values are displayed at the top right of each correlation plot, demonstrating improved structural accuracy with increasing bottleneck dimensions.

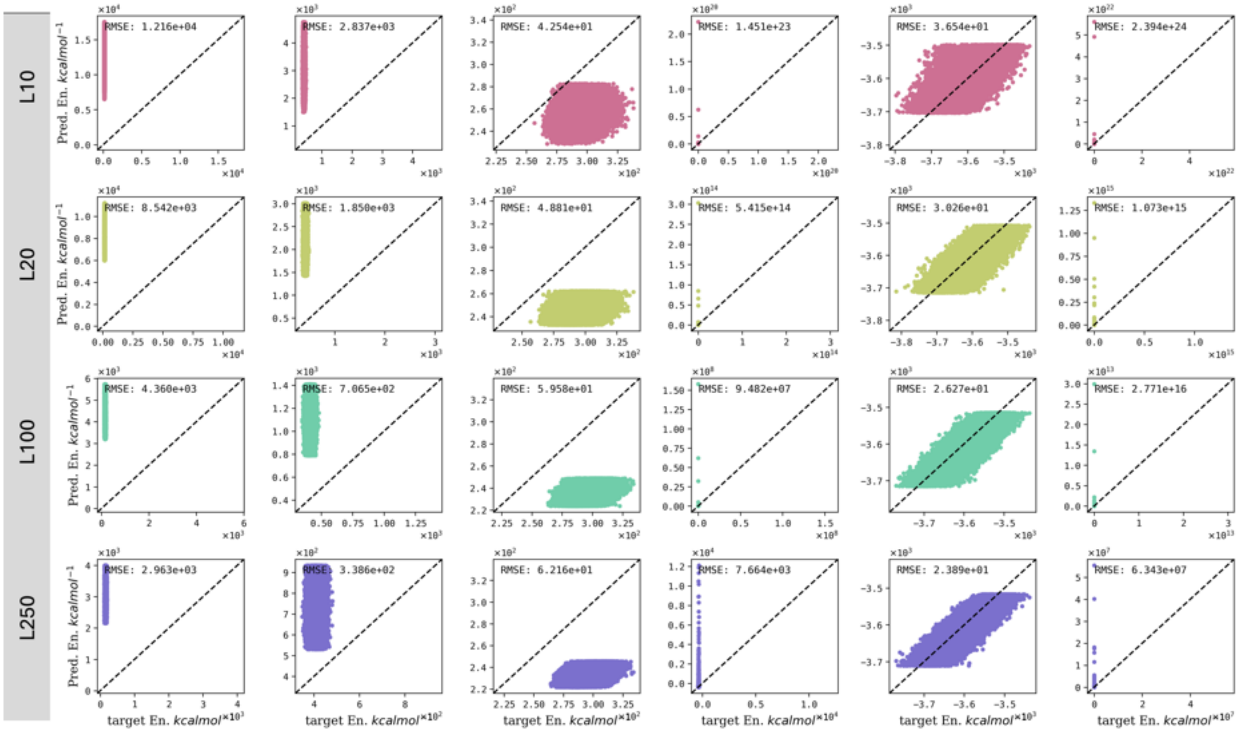


SI Figure 21: Energy component reconstruction analysis across different bottleneck dimensions. Each row represents a distinct latent dimension as labeled (L10: 10 dimensions, L20: 20 dimensions, L100: 100 dimensions, L250: 250 dimensions). Columns show correlation plots of predicted versus target values for bond energy (column 1), angle energy (column 2), dihedral energy (column 3), van der Waals energy (column 4), electrostatic energy (column 5), and total energy (column 6). RMSE values are displayed at the top right of each correlation plot, demonstrating the relationship between bottleneck size and energy reconstruction accuracy across different force field components.
